# Immortalization of human zone I hepatocytes from biliary atresia with CDK4^R24C^, cyclin D1, and TERT for cytochrome P450 induction testing

**DOI:** 10.1101/729525

**Authors:** Manami Nishiwaki, Masashi Toyoda, Yoshie Oishi, Seiichi Ishida, Shin-ichiro Horiuchi, Hatsune Makino, Tohru Kimura, Shin-ichi Ohno, Takashi Ohkura, Shin Enosawa, Hidenori Akutsu, Atsuko Nakazawa, Mureo Kasahara, Tohru Kiyono, Akihiro Umezawa

**Affiliations:** Center for Regenerative Medicine, National Center for Child Health and Development, Tokyo, 157-8535, Japan; Research team for Geriatric Medicine (Vascular Medicine), Tokyo Metropolitan Institute of Gerontology, Tokyo, 173-0015, Japan; Division of Pharmacology, National Institute of Health Sciences, Kanagawa, 210-9501, Japan; Laboratory of Stem Cell Biology, Department of Biosciences, Kitasato University School of Science, Kanagawa 252-0373, Japan; Division for Advanced Medical Sciences, National Center for Child Health and Development, Tokyo, 157-8535, Japan; Saitama Children’s Medical Center, Saitama, 330-8777, Japan; Organ Transplantation Center, National Center for Child Health and Development, Tokyo, 157-8535, Japan; Department of Cell Culture Technology, National Cancer Center Research Institute, Tokyo, 104-0045, Japan; Division of Carcinogenesis and Cancer Prevention, National Cancer Center Research Institute, Tokyo, 104-0045, Japan

**Author notes:** Correspondence should be directed to: Akihiro Umezawa and Tohru Kiyono Akihiro Umezawa, Tohru Kiyono.

**Keywords:** Hepatic stem cell, Toxicology, Drug metabolism, Cytochrome P450, HepaRG, HepG2

## Abstract

**Background:** Hepatocytes are an important tool for in vitro toxicology testing. In addition to primary cultures, a limited number of immortalized cell lines have been developed. We here describe a new cell line, designated as HepaMN, which has been established from a liver associated with biliary atresia.

**Methods:** Hepatocytes were isolated from a liver of 4-year-old girl with biliary atresia and immortalized by inoculation with CSII-CMV-TERT, CSII-CMV-Tet-Off, CSII-TRE-Tight-cyclin D1 and CSII-TRE-Tight-CDK4R24C (mutant CDK4: an INK4a-resistant form of CDK4) lentiviruses at the multiplicity of infection of 3 to 10.

**Results:** HepaMN cells exhibited morphological homogeneity, displaying hepatocyte-like phenotypes. Phenotypic studies in vivo and in vitro revealed that HepaMN cells showed polarized and functional hepatocyte features along with a canalicular cell phenotype under defined conditions, and constitutively expressed albumin and carbamoyl phosphate synthetase I in addition to epithelial markers. Since HepaMN cells are immortal and subcloned, kinetics and expression profiles were independent of population doublings.

**Conclusions:** HepaMN cells showed increased CYP3A4 expression after exposure to rifampicin, implying that their close resemblance to normal human hepatocytes makes them suitable for research applications including drug metabolism studies.

## Background

The liver is essential for maintaining normal physiology and homeostasis and is composed of hepatocytes, endothelial cells, and stellate cells. Among these cells, hepatocytes play a key role in metabolism and detoxification. However, human hepatocytes are difficult to propagate ex vivo due to lack of appropriate cultivation conditions. To solve this issue, hepatocytes have been freshly isolated from livers, generated from hepatomas, induced from pluripotent stem cells such as embryonic stem cells (ESCs) and induced pluripotent stem cells (iPSCs), and converted from other somatic cells such as fibroblasts (1-6). However, a steady supply of hepatocytes dissociated from livers for use as in vitro models cannot be guaranteed because of limited supply and lot-to-lot variations due to genetic and environmental backgrounds. Likewise, iPSC- or ESC-derived hepatocytes may show variation between lots, due to the difficulty of strictly controlling the differentiation process (7). Even when hepatocytes are generated from a bank of undifferentiated iPSCs or ESCs, the direct reprogramming technology requires complex protocols and a relatively long period for complete differentiation, and yields a limited number of mature hepatocytes among a heterogeneous population (8, 9). HepG2, a hepatoblastoma cell line, exhibits hepatocyte-like features with a limited expression of hepatocyte-associate markers such as albumin and cytochrome P450 (CYP) (10). Likewise, HepaRG, a spontaneously immortalized cell line, from hepatocarcinoma of a female patient has more hepatic features compared to HepG2 (11).

Immortalized hepatocytes derived from normal hepatocytes would be ideal to ensure of a steady supply. Immortalized hepatic cell lines have advantages such as formation of large lots, in vitro propagation, genetic and environmental uniformity, and a lack of ethical concerns, compared with cryopreserved isolated hepatocytes and primarily cultivated hepatocytes. From this viewpoint, HepG2 and HepaRG cells have been extensively used for evaluating the toxicity of chemicals and drugs. HepaRG has been used for CYP induction studies, especially induction of CYP3A4, because more than half of all small-molecule drugs commonly used by humans are metabolized by CYP3A4, and inhibition of CYP3A4-mediated metabolism is a common cause of drug-induced liver injury. In this study, we generated an immortalized hepatocyte cell line, HepaMN, from a Japanese patient with biliary atresia. We applied a previously used strategy for immortalization of human keratinocyte or mammary epithelial cells - inactivation of the Rb/p16 pathway and acquisition of telomerase activity (12). HepaMN cells constitutively exhibited a hepatocytic phenotype both in vitro and in vivo, and showed increased CYP3A4 after exposure to rifampicin, implying that HepaMN cells are a suitable tool for pharmaceutical studies.

## Methods

### Viral vector construction and viral transduction

Construction of the lentiviral vector plasmids CSII-CMV-Tet-Off, CSII-TRE-Tight-cyclin D1, and CSII-TRE-Tight-CDK4R24C was previously described (13). In brief, the EF1a promoter in CSII-EF-RfA (a gift from Dr. H. Miyoshi, RIKEN, presently Keio University) was replaced with a tetracycline-inducible promoter, TRE-Tight, from pTRE-Tight (Clontech, Mountain View, CA) to generate CSII-TRE-Tight-RfA. Human cyclin D1, human mutant CDK4 (CDK4^R24C^: an INK4a-resistant form of CDK4), and TERT were inserted into the entry vector via a BP reaction (Invitrogen, Carlsbad, CA). These segments were then recombined with CSII-TRE-Tight-RfA through an LR reaction (Invitrogen) to generate CSII-TRE-Tight-cyclin D1, CSII-TRE-Tight-CDK4R24C, and CSII-TRE-Tight-TERT. The rtTA segment from pTet-Off Advanced (Clontech) was amplified by PCR, recombined with the donor vector pDONR221 via a BP reaction (Invitrogen) to generate pENTR221-Tet-Off, and then recombined with a lentiviral vector, CSII-CMV-RfA, through an LR reaction (Invitrogen) to generate CSII-CMV-Tet-Off. Recombinant lentiviruses with vesicular stomatitis virus G glycoprotein were produced as described previously (14). Primary hepatocytes were inoculated with CSII-CMV-TERT, CSII-CMV-Tet-Off, CSII-TRE-Tight-cyclin D1 and CSII-TRE-Tight-CDK4R24C lentiviruses at the multiplicity of infection of 3 to 10 in the presence of 4 micrograms/mL of polybrene.

### Human Cells

The cells were isolated from a liver of 4-year-old girl with biliary atresia (Hep2018). The hepatocytes were isolated according to the collagenase perfusion method, as described elsewhere (1). The samples were minced into small pieces, washed with HEPES buffer (pH 7.7; 140 mM NaCl/2.68 mM KCl/0.2 mM Na2HPO4/10 mM Hepes), and digested with 0.5 mg/mL collagenase/DMEM (Boehringer Mannheim) diluted in the same buffer supplemented with 0.075% CaCl2 under gentle stirring at 37°C. The cell suspension was washed twice in Hepes buffer and resuspended in a HCM™ BulletKit™ medium (cat. CC-3198, LONZA) supplemented with 10% FBS, hEGF, Transferrin, hydrocortisone, BSA, ascorbic acid, insulin, Fungison, 100 units/ml penicillin, 100 μg/ml streptomycin, 5 μg/ml insulin, and 5 × 10^7^ M hydrocortisone hemisuccinate. Cell suspension containing 2.2 × 10^6^ cells per vial was once frozen for future use. The frozen cells were later thawed and seeded on four 35-mm dishes (BD Falcon 6-well plate) with feeder cells of irradiated human bile duct mesenchymal cells in the ESTEM-MN medium (GlycoTechnica Ltd.) [modified F-medium (DMEM:F12(1:3), 5% FBS, 8.4 ng/ml Cholera Toxin, 30.5 μg/ul Adenine HCl, 10 ng/ml EGF, 0.4 μg/ml Hydrocortisone, 5 μg/ml human insulin, 1% Pen/Strep)] supplemented with 10 uM Y-27632 and 5% conditioned media containing Wnt3a and RSpo1. Y-27632 was supplemented in each passage. Twenty days after plating, several colonies appeared, and nine colonies with hepatocyte-like morphology was picked up by using stainless cloning cylinder with trypsin. The isolated cells were re-plated on a 35-mm dish without any coating feeder layer, and one of them (clone #9) was inoculated with recombinant lentiviruses constitutively expressing TERT and conditionally expressing Cyclin D1 and mutant CDK4 on the same day (13). Cells proliferated constantly after infection, and were frozen at passage 3. These cells were maintained in the ESTEM-MN medium, and the medium was renewed every 2 or 3 days, and were passaged every 1 week (1/5 dilution) by trypsinization.

### Cell culture

HepaMN cells were passaged every 1 week (1/5 dilution) by trypsinization. HepaMN cells were maintained for 1 weeks in the ESTEM-MN medium (GlycoTechnica Ltd.) at 37°C in 5% CO_2_, and the ESTEM-MN medium was renewed every 2 or 3 days.

### Calculation of population doublings

Primary hepatocytes and HepaMN cells were seeded at 0.8 to 1.0 × 10^6 cells/well in a 100-mm dish. When the cells reached subconfluent, both primary cells and HepaMN cells were harvested and the total number of cells in each well was determined using the cell counter. Population doubling (PD) was used as the measure of the cell growth rate. PD was calculated from the formula PD ¼ log2 (A/B), where A is the number of harvested cells and B is the number of plated cells.

### Immunoblotting

Whole-cell protein extracts were used for analysis, and immunoblotting was conducted as described previously (15). Antibodies against CDK4 (clone D9G3E; Cell Signaling Technology, Danvers, MA), CyclinD1 (clone G124-326; BD Biosciences, Franklin Lakes, NJ), GAPDH (clone 6C5, Ambion), p16INK4a (clone G175-405, BD Biosciences) were used as probes, and horseradish peroxidase-conjugated anti-mouse or anti-rabbit immunoglobulins (Jackson Immunoresearch Laboratories, West Grove, PA) were employed as secondary antibodies. The LAS3000 charge-coupled device (CCD) imaging system (Fujifilm Co. Ltd., Tokyo, Japan) was employed for detection of proteins visualized by Lumi-light Plus Western blotting substrate (Roche, Basel, Switzerland).

### Histological analysis of HepaMN

HepaMN cells were harvested with a cell scraper and collected into tubes. The cells were analyzed with an iPGell kit (GenoStaff, Tokyo, Japan).

### Immunocytochemical analysis

HepaMN cells were fixed in 4% paraformaldehyde for 10 min at 4°C. After washing with PBS and treatment with 0.1% Triton-X for 10 min at room temperature, cells were pre-incubated with blocking buffer (1% BSA in PBS) for 30 min at room temperature, and then reacted with primary antibodies in blocking buffer for 12 h at 4°C. Followed by washing with PBS, cells were incubated with secondary antibodies; anti-goat or anti-mouse IgG conjugated with Alexa 488 or 546 (1:1000) (Invitrogen) in blocking buffer for 30 min at room temperature. Then the cells were counterstained with DAPI and mounted.

### Karyotypic analysis

Karyotypic analysis was contracted out at Nihon Gene Research Laboratories Inc. (Sendai, Japan). Metaphase spreads were prepared from cells treated with 100 ng/mL of Colcemid (Karyo Max, Gibco Co. BRL) for 6 h. The cells were fixed with methanol:glacial acetic acid (2:5) three times, and dropped onto glass slides (Nihon Gene Research Laboratories Inc.). Chromosome spreads were Giemsa banded and photographed. A minimum of 10 metaphase spreads were analyzed for each sample, and karyotyped using a chromosome imaging analyzer system (Applied Spectral Imaging, Carlsbad, CA).

### Quantitative RT-PCR

RNA was extracted from cells using the ISOGEN (NIPPON GENE). An aliquot of total RNA was reverse transcribed using an oligo (dT) primer (SuperScript TM III First-Strand Synthesis System, Invitrogen). For the thermal cycle reactions, the cDNA template was amplified (QuantStudio TM 12K Flex Real-Time PCR System) with gene-specific primer sets (Supplemental Table 1) using the Platinum Quantitative PCR SuperMix-UDG with ROX (11743-100, Invitrogen) under the following reaction conditions: 40 cycles of PCR (95°C for 15 s and 60°C for 1 min) after an initial denaturation (50°C for 2 min and 95°C for 2 min). Fluorescence was monitored during every PCR cycle at the annealing step. The authenticity and size of the PCR products were confirmed using a melting curve analysis (using software provided by Applied Biosystems) and gel analysis. mRNA levels were normalized using ubiquitin or GAPDH as a housekeeping gene.

### Gene chip analysis

High quality total RNA was isolated from each cell using Trizol® following the manufacturer’s instructions (Invitrogen, USA). Genomic DNA was eliminated by treatment with DNAse I for 20 min at RT using DNAse IH (Invitrogen, USA). RNA concentration was measured using a Nanodrop ND-1000 spectrophotometer (NanoDrop Technologies, Wilmington, Delaware USA). Purity and integrity of total RNA was determined by 260/280 nm ratio and checked by electrophoresis in Bioanalyzer RNA6000 Nano. About 100 ng of total RNA was used to produce Cyanine 3-CTP labeled cRNA using the Low Input Quick Amp Labeling Kit, One-Color (Agilent Technologies) according to the manufacturer’s instructions. Following ‘One-Color Microarray Based Gene Expression Analysis’ protocol version 6.0 (Agilent Technologies), 2 μg of labeled cRNA was hybridized with a human gene expression microarray 60K (Agilent Technologies, Santa Clara CA, USA). The microarray workflow quality control was implemented using the Agilent Spike-In Kit which consisted of a set of 10 positive control transcripts optimized to anneal to complementary probes on the microarray with minimal self-hybridization or cross-hybridization. The concentrated Agilent One-Color RNA Spike-In mix stock was diluted in the buffer provided by the kit and mixed with the RNA samples prior to the amplification and labeling process to achieve the relative amounts recommended by the manufacturer.

For hybridization, Agilent gene expression microarray 60K slide (Design ID: 72363. SurePrint G3 Human Gene Expression 8 × 60K Microarray Kit, Agilent Technologies) were used. Slides were scanned in an Agilent C Scanner according to the manufacturer’s protocol. Signal data were collected with dedicated Agilent Feature Extraction Software (v 11.5.1). Agilent Processed Signals were processed using GeneSpring software (Agilent Technologies).

### Principal component analysis (PCA)

To analyze the expression data of genes in an unsupervised manner with a gene chip array, we used principal component analysis (16, 17). PCA, a multivariate analysis technique which finds major patterns in data variability, was performed to categorized HepaMN cells into each stage of ESCs undergoing hepatic differentiation. The differentiation protocol was previously reported in detail (18). For ESC-derived hepatocytes, Sees2 was maintained in the XF32 medium [85% Knockout DMEM, 15% Knockout Serum Replacement XF CTS (XF-KSR), 2 mM GlutaMAX-I, 0.1 mM NEAA, Pen-Strep, 50 µg/mL L-ascorbic acid 2-phosphate, 10 ng/mL heregulin-1β (recombinant human NRG-β 1/HRG-β 1 EGF domain), 200 ng/mL recombinant human IGF-1, and 20 ng/mL human bFGF. To generate embryoid bodies (EBs), ESCs (1 × 10^4^/well) were dissociated into single cells with accutase after exposure to the rock inhibitor (Y-27632), and cultivated in the 96-well plates in the EB medium [76% Knockout DMEM, 20% Knockout Serum Replacement, 2 mM GlutaMAX-I, 0.1 mM NEAA, Pen-Strep, and 50 µg/mL L-ascorbic acid 2-phosphate for 10 days. The EBs were transferred to the 24-well plates, and cultivated in the XF32 medium for 14 to 35 days.

### In vivo analysis

The operation protocols were approved by the Laboratory Animal Care and the Use Committee of the National Research Institute for Child and Health Development, Tokyo. HepaMN cells were harvested with a cell scraper, collected into tubes, centrifuged, and resuspended in PBS (200 μl). The same volume (200 μl) of Matrigel was added to the cell suspension. The cells (> 1 × 10^7^) were subcutaneously inoculated into severe combined immunodeficiency (SCID) mice (CREA, Tokyo, Japan). After 9 days, the resulting tumors were dissected and fixed with PBS containing 4% paraformaldehyde. Paraffin-embedded tissue was sliced and stained with hematoxylin and eosin (HE). HepaMN cells were also implanted into spleen in immunodeficient SCID mice. HepaMN cells were harvested with a cell scraper, collected into tubes, centrifuged, and resuspended in PBS (50 μl). The cells (> 1 × 10^6^) were intrasplenically implanted. After 7 days, the spleens were dissected and fixed with PBS containing 4% paraformaldehyde for further histological analysis.

### Immunohistochemistry

Subcutaneous nodules on mouse backs were fixed in 20% formalin and embedded in paraffin. Cut paraffin sections were deparaffinized, dehydrated, and treated with 2% proteinase K (Dako) in Tris-HCl buffer solution (pH 7.5) for 5 min at room temperature, or heated in ChemMate Target Retried Solution (Dako) for 5-20 min in a high-pressure steam sterilizer for epitope unmasking. After washing with distilled water, samples were placed in 1% hydrogen peroxide/methanol for 15 min to block endogenous peroxidase. The sections were then incubated at room temperature for 60 min in primary antibodies diluted with antibody diluent (Dako). The following primary antibodies against various human differentiation antigens were used: vimentin (V9, M0725, Dako, Glostrup, Denmark), albumin (Dako) and AE1/AE3 (712811, NICHIREI). Then, they were washed three times with 0.01 M Tris buffered saline (TBS) solution (pH 7.4) and incubated with goat anti-mouse or anti-rabbit immunoglobulin labeled with dextran molecules and horseradish peroxidase (EnVision, Dako) at room temperature for 30 min. After three times washes with TBS, they were incubated in 3,3’-diaminobenzidine in substrate-chromogen solution (Dako) for 5-10 min. Negative controls were performed by omitting the primary antibody. The sections were counterstained with hematoxylin.

### CYP3A4 induction test

HepaMN cells were cultivated in the 24-well plates, and was treated with 20 μM rifampicin or 125 μM omeprazole for 24 h at a cell density of 80%. Controls were treated with DMSO (final concentration 0.1%). Controls were treated with DMSO.

### CYP3A4 Activity and Chromatographic Conditions

The reaction mixtures [0.05 M potassium phosphate buffer (pH 7.4), 2 mM NADPH (Trevigen Inc., MD), 20 mM 7-benzyloxy-4-trifluoromethylcoumarin (BFC, Sigma-Aldrich, MD) and cell homogenate (20 – 100 mg protein) in a final volume of 200 ml] were incubated at 37°C for 1 h (19). Reactions were terminated at 95°C for 2 min, followed by centrifugation at 10,000 x g for 5 min. For the inhibition assay by ketoconazole (KTZ, Cayman Chemical Co., Michigan, USA) as inhibitor of human CYP3A isoform, KTZ in 50% methanol solution, yielding final concentrations 10 mM in the incubation mixture, was added to the incubation tubes (20). A Coomassie Plus (Bradford) Protein Assay kit (Thermo Fisher Scientific, MA, USA) was used to determine protein concentrations with bovine serum albumin as the standard.

Quantitative analysis of 7-Hydroxy-4-trifluoromethylcoumarin (HFC) produced by CYP3A4 was performed by reverse-phase high performance liquid chromatography system (RP-HPLC) equipped with a Cosmosil 5C_18_-AR-II column (4.6 I.D. x 150 mm, Nacalai Tesque, Kyoto, Japan). RP-HPLC analysis was carried out in isocratic conditions using a reciprocating dual pump KP-22 (FLOM Inc., Tokyo, Japan) and a Rheodyne manual injector with a loop size of 100 μl. Running conditions included: injection volume, 50 μl; mobile phase, methanol: 0.02 M phosphate buffer (pH7.4) (3: 4 v/v); flow rate, 0.2 ml/min; and fluorescence detection at an excitation wavelength of 410 nm and an emission wavelength of 510 nm using a Multi *λ* Fluorescence Detector 2475 (Waters Co., Milford, MA). Calibration curves known amounts of HFC (Sigma-Aldrich, MD) were added to the mixture of buffer– methanol (1:1). The linear concentration range was 1 to 100 nmol/ml for HFC.

### Statistical analysis

Statistical analysis was performed using the unpaired two-tailed Student’s t test.

## Results

### Generation of HepaMN cells

Cells were isolated from a 4-year-old patient with biliary atresia (Figure 1A-D). These cells had a hepatocyte-like morphology after primary culture, and were immortalized by the introduction of CDK4R24C, cyclin D1, and TERT. Immortalized hepatocytes, designated as HepaMN cells, expressed CDK4R24C, cyclin D1, and TERT and resembled primary cells in morphology (Figure 1E, F). HepaMN cells appeared as a homogeneous cell population with an epithelial phenotype showing no regular structural organization. Even after reaching confluence, the cells retained the appearance of hepatocyte-like cells. Morphological characteristics of HepaMN cells did not significantly change even at later passages.

**Figure 1.**
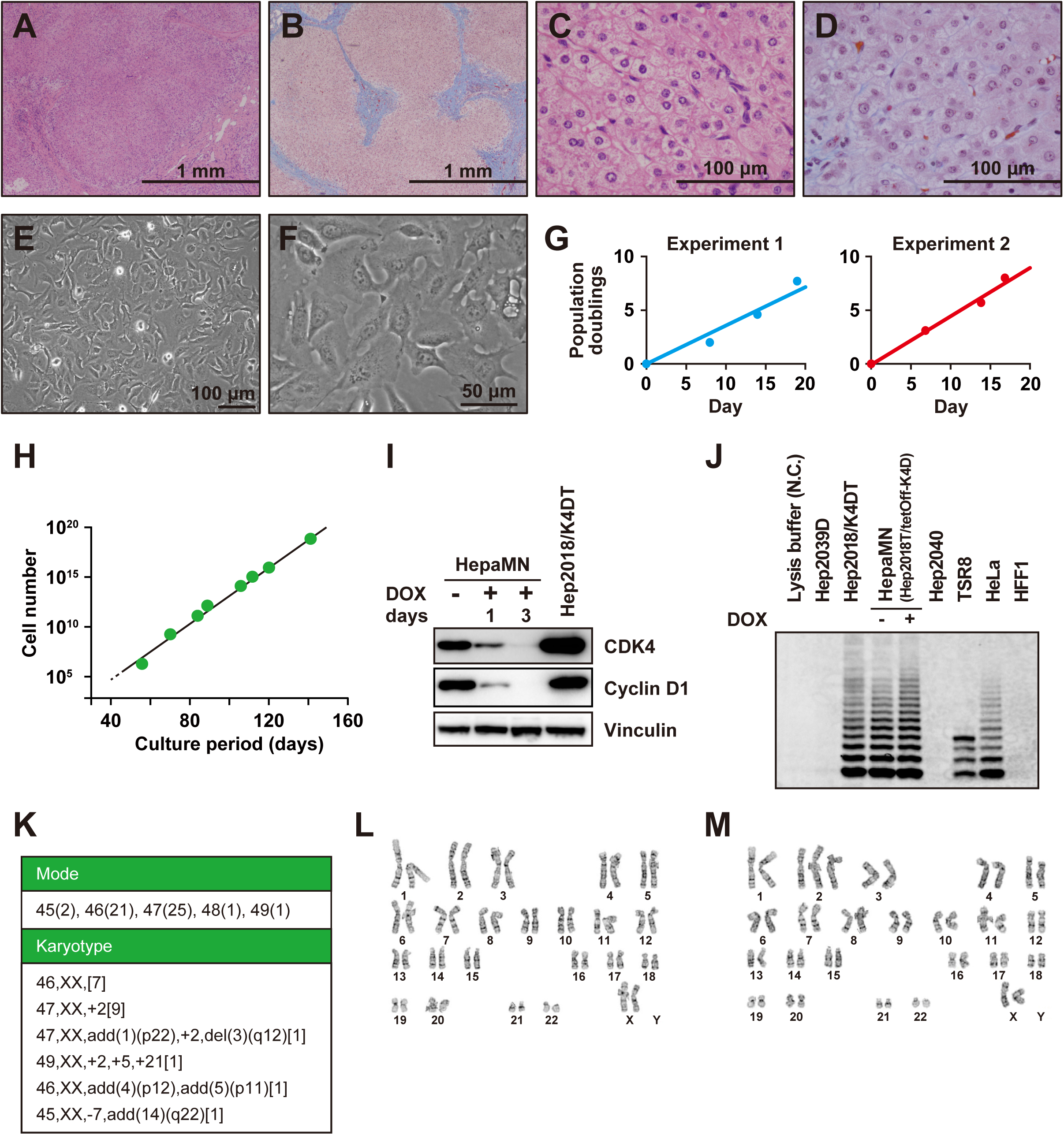
Establishment of HepaMN cells. A. Histology of the liver from which the hepatocytes were isolated. HE stain. B. Masson-Trichrome stain of the liver. C. High power view of panel A. D. High power view of panel B. E. Phase contrast photomicrograph of HepaMN cells at a subconfluence. F. High power view of panel E. G. Growth curve of HepaMN cells in independent duplicate experiments (Experiment 1: left, Experiment 2: right). HepaMN cells were appropriately passaged every 1 week (1/5 dilution) by trypsinization. H. in vitro growth of HepaMN cells. HepaMN cells proliferated up to 10^18^ cells for more than 140 days. I. Western blot analysis of cell cycle-associated protein levels in HepaMN cells. Expression of CDK4 and CYCLIN D1 in HepaMN cells grown in the absence or presence of doxycycline (DOX) for 1 and 3 days were analyzed. VINCULIN was used as a loading control. Hep2018/K4DT cells were used as a positive control for expression of CDK4 and CYCLIN D1. J. Telomerase activity of HepaMN cells. Telomerase activity is revealed by the characteristic six-base pair ladder of bands. Telomerase activity was detected in Hep2018/K4DT and HepaMN cells, but not in non-transduced cells, i.e. Hep2039D and Hep2040 cells. TSR8 and HeLa cells serve a positive control with telomerase activity and human foreskin fibroblasts (HFF1) serve a negative control without telomerase activity. N.C.: Negative control. K. Karyotypic analyses of HepaMN cells at Passage 16. For the mode analysis, 50 cells were analyzed and the number of cells with the indicated chromosomal number was shown in parenthesis. For karyotypic analysis, 20 cells were analyzed and the number of cells with the indicated karyotype was shown in brackets. L. Karyogram of a HepaMN metaphase at Passage 16 with the G band method. Normal chromosomes are seen. M. Karyogram of a HepaMN metaphase at Passage 16 with the G band method indicating trisomy of chromosome 2.

HepaMN cells reproducibly proliferated more rapidly than the primary cells as determined by population doubling values (Figure 1G). HepaMN cells continued to expand, and neither cessation of cell proliferation, such as senescence nor apoptosis/cell death, was detected during cultivation through 21 passages (141 days), implying that the hepatocytes were immortalized by the introduction of CDK4^R24C^, cyclin D1, and TERT genes (Figure 1H). We performed immunoblot analysis of CDK4 and cyclin D1 to confirm the expression of the transgenes (Figure 1I). We found that CDK4 and cyclin D1 were highly expressed, and suppressed by doxycycline treatment. To determine TERT, we performed a TRAP assay in a series of hepatocytes infected with the TERT gene (Figure 1J). The results clearly show that the TERT infection resulted in the induction of enzyme activity. We also performed karyotypic analysis of HepaMN cells. HepaMN cells had a normal karyotype (7 out of 20 metaphases) and chromosome 2 trisomy (9 out of 20 metaphases) at passage 16 (Figure 1K, L, M).

### Characterization of HepaMN cells in vitro

Immunocytochemistry revealed that HepaMN cells were positive for albumin and epithelial keratins (Figure 2A-D). Interestingly, HepaMN cells also positively stained for the mesenchymal marker vimentin, and cells positive for both keratins (AE1/AE3) and vimentin were observed (Figure 2E). Vimentin, a marker for a mesenchymal phenotype, is induced in cells undergoing epithelial-to-mesenchymal transition under certain conditions, such as development and oncogenesis, and indeed hepatocytes without introduction of the genes for immortalization were negative for vimentin (Supplemental Figure 1). HepaMN cells are positive after the immortalization process. The other immortalized hepatocytes analyzed (immortalized cells from Hep2040, Hep2023D, Hep2020, and Hep2013) are all positive for vimentin. The paraffin-embedded HepaMN cells were again positive for vimentin, and contained glycogen in the cytoplasm by PAS stain (Figure 2F-J). Electron microscopy studies of HepaMN hepatocytes showed numerous glycogen granules in the cytoplasm (Figure 2K, L). HepaMN hepatocytes exhibited desmosomes between adjacent cells, with structures resembling bile canaliculi, delineated by typical junctional complexes and presenting several microvilli (Figure 2M, N). HepaMN cells contained numerous subcellular organelles, such as mitochondria, peroxisomes, lysosomes and endoplasmic reticulum (Figure 2O, P).

**Figure 2.**
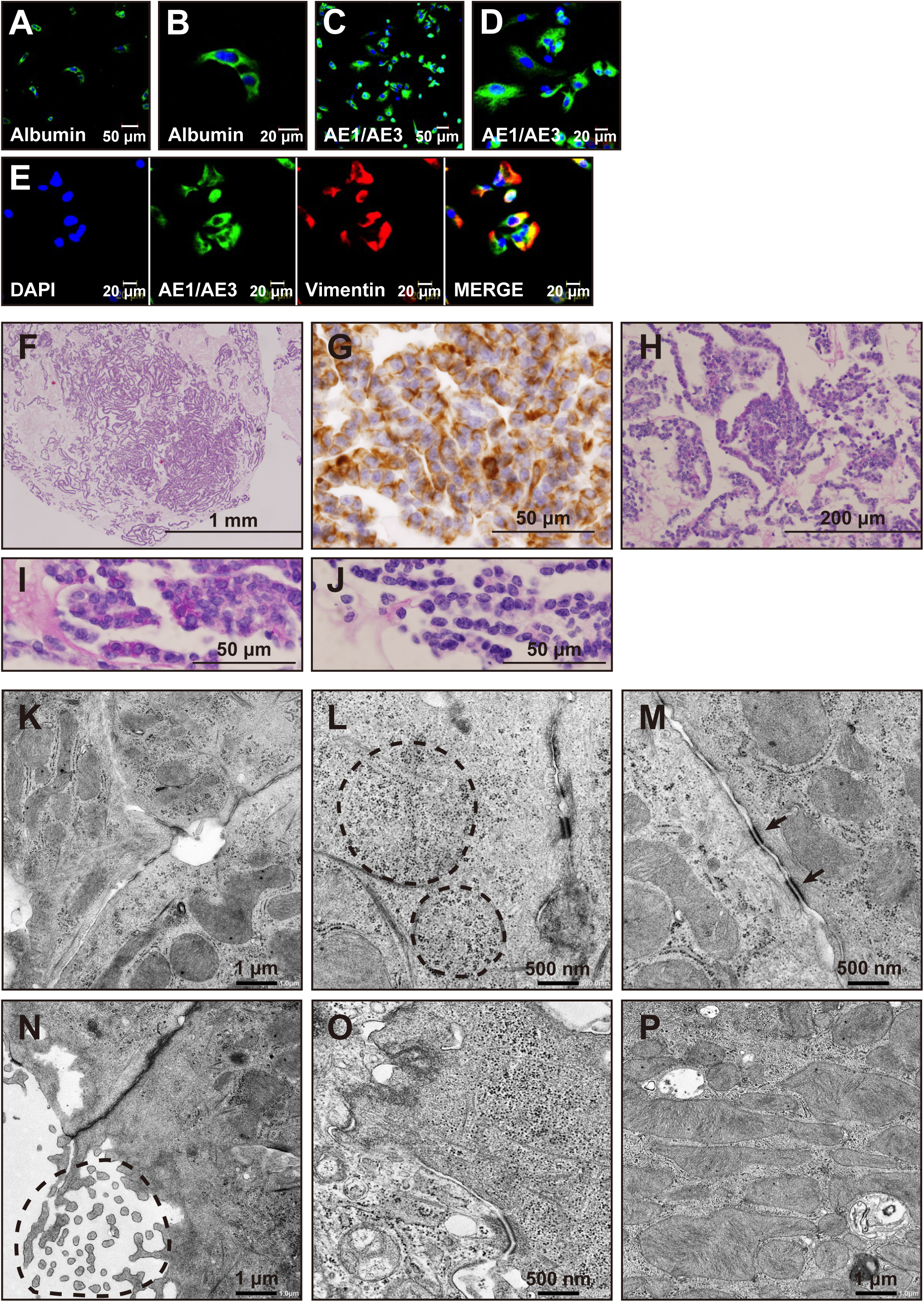
Characterization of HepaMN cells. A. Immunocytochemistry of HepaMN cells with an antibody to albumin. B. High power view of panel H. C. Immunocytochemistry of HepaMN cells with AE1/3. D. High power view of panel J. E. Immunocytochemistry of HepaMN cells with AE1/3 (green) and vimentin (red). Nuclei were stained with DAPI (blue). F. HepaMN cells in iPGell. HE stain. G. Immunocytochemistry of HepaMN cells in iPGell with the antibody to vimentin. H. Glycogen accumulation in HepaMN cells. Periodic acid–Schiff (PAS) stain. I. High power view of panel H. J. PAS stain with diastase digestion of panel I. K. Electron microscopy study of HepaMN cells. HepaMN cells exhibited hepatocyte-like morphology evidenced contacting cells tightly attached by desmosomes, with structures identical to bile canaliculi. L. Glycogen β-particles in the HepaMN cytoplasm. HepaMN cells accumulated abundant glycogen particles. M. Desmosomes between HepaMN cells. N. Microvilli on the HepaMN membrane. HepaMN cells contained microvilli which project into the lumen from each HepaMN cell. O. Endoplasmic reticulum in the HepaMN cytoplasm. P. Abundant mitochondria in the HepaMN cytoplasm.

### Hepatocyte-associated gene expression in HepaMN cells

HepaMN cells were monitored by analyzing a representative panel of liver-specific genes, all normally expressed in adult hepatocytes (Figure 3A-I, Supplemental Table 1). RNA levels were examined in HepaMN cells and compared with the differentiated human hepatoblastoma-derived HepaRG cell line. HepaMN cells expressed the albumin gene at a higher level than HepaRG, but expressed the other liver-specific genes at low or undetectable levels.

**Figure 3.**
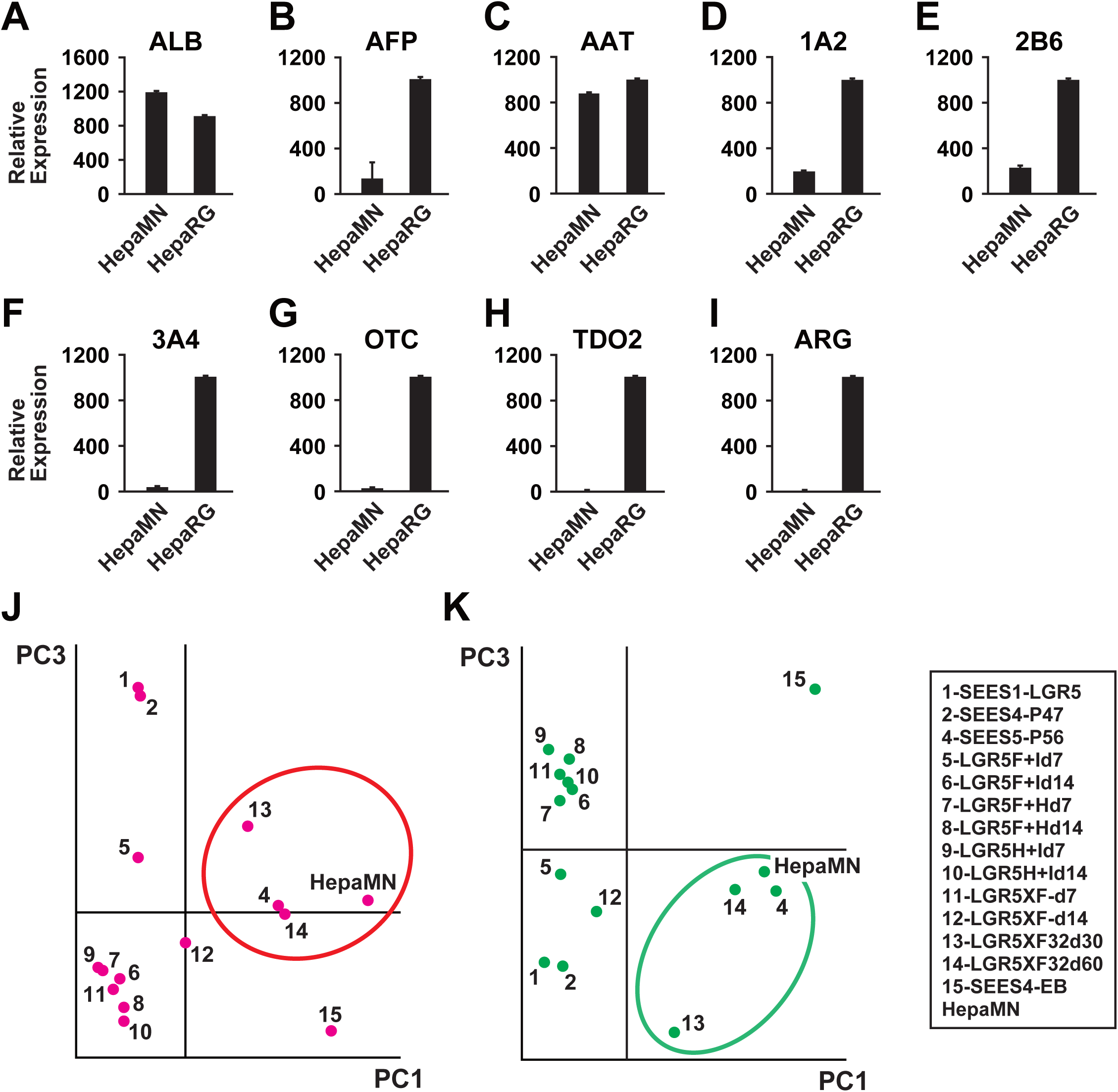
Liver-specific gene expression in HepaMN cells. A. Albumin (ALB). Quantitative RT-PCR analysis was performed on HepaMN cells. mRNA levels were normalized using ubiquitin as a housekeeping gene. B. α-fetoprotein (AFP). C. α-antitrypsin (AAT). D. CYP1A2 (1A2). E. CYP2B6 (2B6). F. CYP3A4 (3A4). G. Ornithine transcarbamylase (OTC). H. Tryptophan 2,3-dioxygenase (TDO2). I. Arginase (ARG). J. Principal component analysis of “Liver Development (Supplemental Table 2A)”-associated gene expression on HepaMN cells, undifferentiated ESCs (1: SEES1-LGR5, 2: SEES4 at Passage 47), and human ESC-derived endodermal cells [4 : SEES5-P56 (SEES5 endodermal cells at day 50), 5: LGR5F+I d7 (SEES1-LGR5 endodermal cells at day 7), 6: LGR5F+I d14 (SEES1 endodermal cells at day 14), 7: LGR5F+H d7 (SEES1 endodermal cells at day 7), 8: LGR5F+H d14 (SEES1 endodermal cells at day 14), 9: LGR5H+I d7 (SEES1 endodermal cells at day 7), 10: LGR5H+I d14 (SEES1 endodermal cells at day 7), 11: LGR5XF-d7 (SEES1 endodermal cells at day 7), 12: LGR5XF-d14 (SEES1 endodermal cells at day 14), 13: LGR5XF32 d30 (SEES1 endodermal cells at day 30), 14: LGR5XF32 d60 (SEES1 endodermal cells at day 60), 15: SEES4-EB (SEES4 embryoid body at day 14] are shown. The detailed protocol and estimated differentiation stage are detailed in Supplemental Table 5. K. Principal component analysis of “Mature Hepatocyte (Supplemental Table 2B)”-associated gene expression on HepaMN cells and human ESC-derived hepatocytes are shown.

### Global outlook by principal component analysis (PCA)

To clarify the differentiated state of HepaMN, we compared the gene expression levels in HepaMN cells and hepatocyte-differentiated ESCs using Agilent Technologies GeneChip oligonucleotide arrays. LGR5XF32d30 and LGR5XF32d60 used in this study are human ESCs that were differentiated into hepatocytes for 30 and 60 days, respectively. We then performed PCA from the expression data of liver-associated genes, i.e. developing and mature hepatocyte markers (Supplemental Table 2). PCA based on liver-associated genes demonstrated that HepaMN cells are grouped into the category that includes embryonic stem cell-derived terminally differentiated hepatocytes (circled in red and green in Figure 3J and 3K, respectively). The genes on the PC1 axis, based on developing and mature hepatocyte markers, are listed with correlations in Table 1A and B, respectively. These similarities led us to hypothesize that HepaMN cells maintain a specific differentiated state that strongly resembles hepatocytes.

**Table 1.** A list of principal components (PC)

| Probe ID | logChange | Correlation | GeneSymbol | Description |
| --- | --- | --- | --- | --- |
| A_24_P73577 | 2.30 | 0.94 | ALDH1A2 | aldehyde dehydrogenase 1 family, member A2 (ALDH1A2) |
| A_23_P150768 | 3.46 | 0.93 | SLCO2B1 | solute carrier organic anion transporter family, member 2B1 (SLCO2B1) |
| A_32_P178800 | 2.25 | 0.94 | ITGA2 | integrin, alpha 2 (CD49B, alpha 2 subunit of VLA-2 receptor) (ITGA2) |
| A_23_P126266 | 2.05 | 0.87 | HLX1 | H2.0-like homeobox 1 (Drosophila) (HLX1) |
| A_24_P34155 | 2.10 | 0.93 | ENST00000358356 | Runt-related transcription factor 1 |
| A_23_P47034 | 2.19 | 0.79 | HHEX | hematopoietically expressed homeobox (HHEX) |
| A_24_P96403 | 2.24 | 0.94 | RUNX1 | runt-related transcription factor 1 (acute myeloid leukemia 1; aml1 oncogene) (RUNX1) |
| A_23_P406782 | 2.57 | 0.93 | HPN | hepsin (transmembrane protease, serine 1) (HPN) |
| A_24_P157926 | 1.31 | 0.90 | TNFAIP3 | tumor necrosis factor, alpha-induced protein 3 (TNFAIP3) |
| A_23_P215790 | 1.34 | 0.89 | EGFR | epidermal growth factor receptor (erythroblastic leukemia viral (v-erb-b) oncogene homolog, avian) (EGFR) |
| A_23_P163402 | 1.73 | 0.86 | CYP1A1 | cytochrome P450, family 1, subfamily A, polypeptide 1 (CYP1A1) |
| A_23_P206733 | 0.83 | 0.87 | CES1 | carboxylesterase 1 (monocyte/macrophage serine esterase 1) (CES1) |
| A_23_P207842 | 0.61 | 0.92 | RARA | retinoic acid receptor, alpha (RARA) |
| A_23_P104689 | 0.38 | 0.74 | RELA | v-rel reticuloendotheliosis viral oncogene homolog A, nuclear factor of kappa light polypeptide gene enhancer in B-cells |
| A_23_P327910 | -1.56 | -0.70 | ZIC3 | Zic family member 3 heterotaxy 1 (odd-paired homolog, Drosophila) (ZIC3) |
| A_23_P24515 | -0.56 | -0.80 | ACAT1 | acetyl-Coenzyme A acetyltransferase 1 |
| A_23_P214977 | -0.43 | -0.74 | SEC63 | SEC63 homolog (S. cerevisiae) (SEC63) |
| A_23_P54556 | -0.46 | -0.74 | MKL2 | MKL/myocardin-like 2 (MKL2) |
| A_32_P210202 | -0.76 | -0.90 | E2F7 | E2F transcription factor 7 (E2F7) |
| A_24_P500891 | -0.38 | -0.80 | AK2 | adenylate kinase 2 (AK2) |
| A_23_P9056 | -0.31 | -0.74 | RB1CC1 | RB1-inducible coiled-coil 1 (RB1CC1) |

**B. PC1 based on mature hepatocyte-associated gene expression**
| Probe ID | logChange | Correlation | GeneSymbol | Description |
| --- | --- | --- | --- | --- |
| A_23_P12767 | 3.82 | 0.97 | CYP2C9 | cytochrome P450, family 2, subfamily C, polypeptide 9 (CYP2C9) |
| A_23_P8801 | 3.54 | 0.99 | CYP3A5 | cytochrome P450, family 3, subfamily A, polypeptide 5 (CYP3A5) |
| A_23_P108280 | 2.25 | 0.88 | CYP4F12 | cytochrome P450, family 4, subfamily F, polypeptide 12 (CYP4F12) |
| A_23_P52480 | 2.65 | 0.87 | CYP2C18 | cytochrome P450, family 2, subfamily C, polypeptide 18 (CYP2C18) |
| A_32_P178800 | 2.04 | 0.91 | ITGA2 | integrin, alpha 2 (CD49B, alpha 2 subunit of VLA-2 receptor) (ITGA2) |
| A_23_P50710 | 1.72 | 0.74 | CYP4F2 | cytochrome P450, family 4, subfamily F, polypeptide 2 (CYP4F2) |
| A_23_P358917 | 2.78 | 0.99 | CYP3A7 | cytochrome P450, family 3, subfamily A, polypeptide 7 (CYP3A7) |
| A_23_P257834 | 2.93 | 0.84 | ALB | albumin (ALB) |
| A_23_P406782 | 2.39 | 0.92 | HPN | hepsin (transmembrane protease, serine 1) (HPN) |
| A_24_P945228 | 1.29 | 0.78 | CYP4V2 | cytochrome P450, family 4, subfamily V, polypeptide 2 (CYP4V2) |
| A_23_P131060 | 1.41 | 0.84 | CYP4F8 | cytochrome P450, family 4, subfamily F, polypeptide 8 (CYP4F8) |
| A_23_P208373 | 1.25 | 0.90 | CYP2B6 | cytochrome P450, family 2, subfamily B, polypeptide 6 (CYP2B6) |
| A_24_P157926 | 1.32 | 0.97 | TNFAIP3 | tumor necrosis factor, alpha-induced protein 3 (TNFAIP3) |
| A_24_P339514 | 2.08 | 0.89 | CYP2B6 | cytochrome P450, family 2, subfamily B, polypeptide 6 (CYP2B6) |
| A_23_P163402 | 1.53 | 0.81 | CYP1A1 | cytochrome P450, family 1, subfamily A, polypeptide 1 (CYP1A1) |
| A_23_P143734 | 0.82 | 0.71 | CYP2D6 | cytochrome P450, family 2, subfamily D, polypeptide 6 (CYP2D6) |
| A_23_P155123 | 0.78 | 0.73 | CYP2D6 | cytochrome P450, family 2, subfamily D, polypeptide 6 (CYP2D6) |
| A_23_P103486 | 1.29 | 0.82 | CYP2J2 | cytochrome P450, family 2, subfamily J, polypeptide 2 (CYP2J2) |
| A_32_P210202 | -0.75 | -0.95 | E2F7 | E2F transcription factor 7 (E2F7) |
| A_23_P160742 | -0.64 | -0.79 | GLMN | glomulin, FKBP associated protein (GLMN) |

### In vivo analysis of HepaMN cells

To address whether HepaMN cells exhibit hepatocytic phenotypes in vivo, they were injected into subcutaneous tissue (1.0 to 5.0 × 10^7^ cells/site) of immunodeficient SCID mice (Figure 4A, B). Twenty independent experiments revealed that HepaMN cells reproducibly showed hepatocytic morphology with canalicular structure in vivo nine days after implantation (Figure 4C, D). Mock surgery with injection of PBS alone did not show the presence of hepatocyte-like cells at the injection sites. Immunohistochemical analysis showed that HepaMN cells were positive for Hep1 (CPS1), albumin, and AE1/3, but negative for α-fetoprotein (AFP) (Figure 4E-L). HepaMN cells were also immunohistochemically analyzed in the spleen seven days after implantation (Figure 4M-V). Again, HepaMN cells were positive for Hep1, albumin, and AE1/3. HepaMN cells also stained positive for MRP2 at the cell membrane and CK19 in the cytoplasm. Nor did HepaMN exhibit cell proliferation histopathologically and produce obvious tumors macroscopically in the subcutaneous tissue of the SCID mice during the monitoring period (>30 days), implying that HepaMN cells do not have an oncogenic potential.

**Figure 4.**
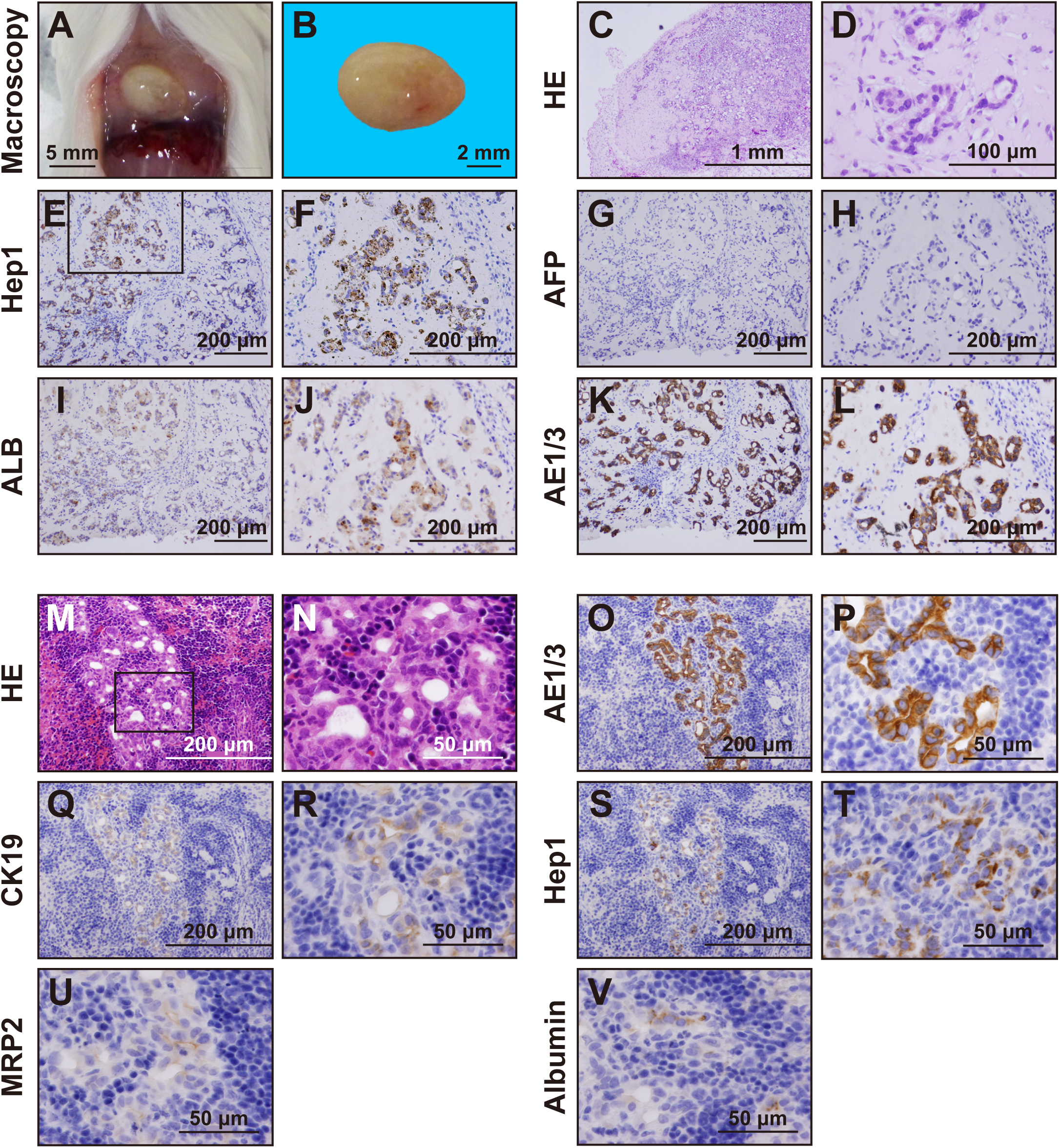
Hepatic characterization of HepaMN cells in vivo. A. Macroscopic view of HepaMN cell-implanted site. HepaMN cells were mixed with MatriGel, and then implanted into the subcutaneous tissue of SCID mice. The HepaMN cell-generated mass was resected at 9 days after the implantation. Bar: 5 mm. B. Macroscopic view of the mass by implantation of HepaMN cells. Bar: 2 mm. C. Histology of the implanted HepaMN cells. HE stain. D High power view of panel C. E. Immunohistochemical analysis of HepaMN cells with an antibody to CPS1 (Hep1). F. High power view of panel E. G. Immunohistochemical analysis of HepaMN cells with an antibody to AFP. H. High power view of panel G. I. Immunohistochemical analysis of HepaMN cells with an antibody to ALB. J. High power view of panel I. K. Immunohistochemical analysis of HepaMN cells with AE1/3. L. High power view of panel K. M. Implantation of HepaMN cells into the spleen. The spleen was resected at 7 days after the implantation. HE stain. N. High power view of panel M. O. Immunohistochemical analysis of HepaMN cells with AE1/3. P. High power view of panel O. Q. Immunohistochemical analysis of HepaMN cells with an antibody to cytokeratin 19 (CK19). R. High power view of panel Q. S. Immunohistochemical analysis of HepaMN cells with an antibody to CPS1 (Hep1). T. High power view of panel S. U. Immunohistochemical analysis of HepaMN cells with MRP2. V. Immunohistochemical analysis of HepaMN cells with albumin (ALB).

### Measurement of cytochrome P450 (CYP) mRNA induction after drug treatment

In order to examine HepaMN cells for CYP induction, HepaMN cells were exposed to omeprazole, phenobarbital, or rifampicin (Figure 5A). CYP3A4 and CYP1A2 were induced with exposure to rifampicin 8.5- and 2.1-fold on average, respectively, while CYP2B6 did not show any induction. CYP1A1 and CYP1A2 were induced with exposure to omeprazole or β-naphthoflavone (Supplemental Figure 2). We further analyzed CYP3A4 metabolism enzyme activity quantitatively in HepaMN cells at sub-confluence and compared the results with the activity observed in HepaRG cells (Figure 5B, C). CYP3A4 activity was 38.6 pmol/min/mg protein, and the CYP3A4 value was lower in HepaMN cells than in HepaRG cells. The CYP3A4 activity in HepaMN cells was specifically inhibited by the specific CYP3A4 inhibitor ketoconazole. Expression levels of albumin and AFP remained unchanged (Figure 5D, E).

**Figure 5.**
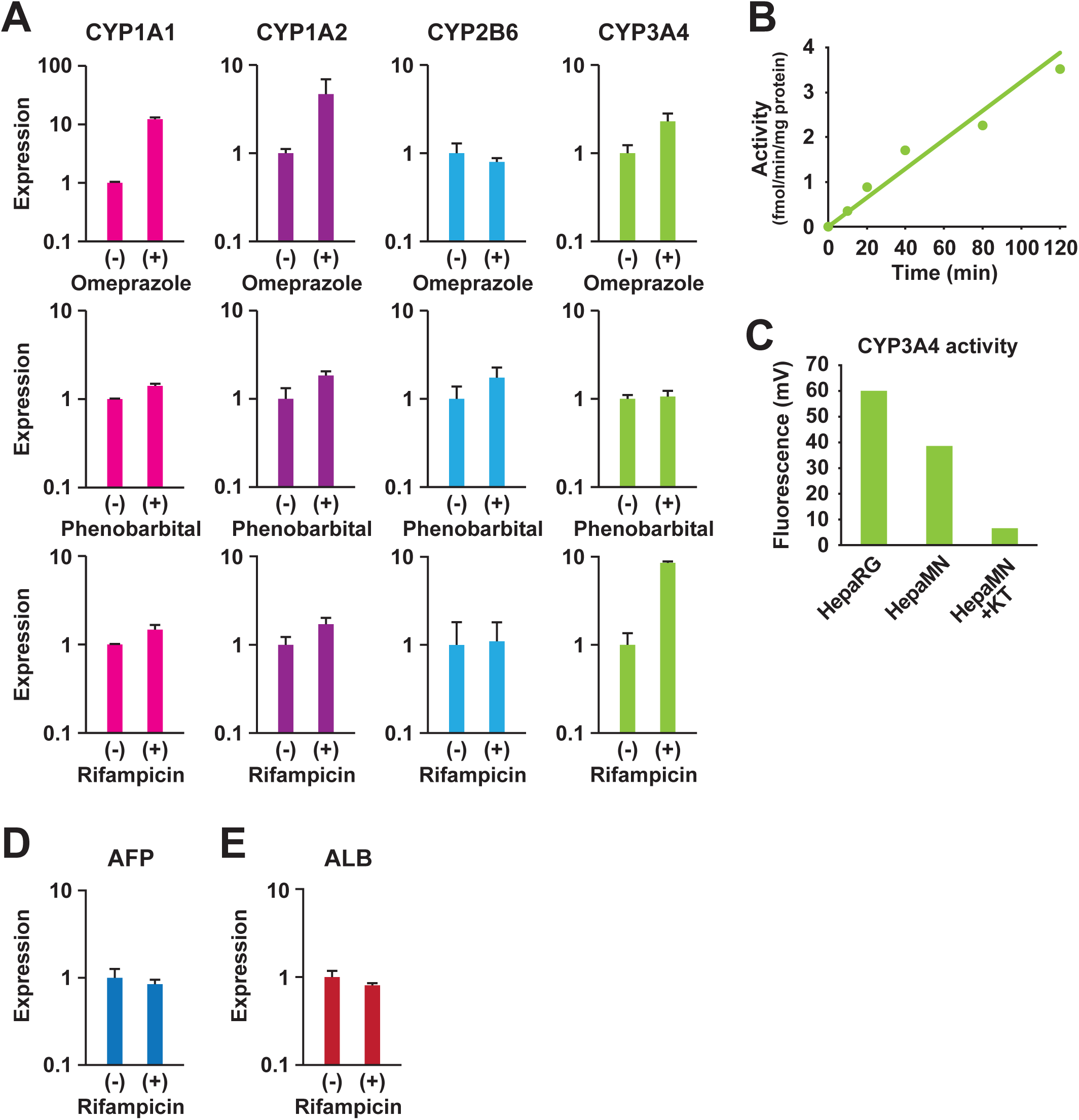
Cytochrome P450 gene induction test in HepaMN cells. A. Quantitative RT-PCR analysis of the genes for CYP1A1, CYP1A2, CYP2B6 and CYP3A4 was performed on HepaMN cells with exposure to 50 μM omeprazole for 24 h, 500 μM phenobarbital for 48 h, or 20 μM rifampicin for 48 h. mRNA levels were normalized using ubiquitin as a housekeeping gene. B. CYP3A4 enzymatic activity in HepaRG and HepaMN cells were measured by using 7-Benzyloxy-4-trifluoromethylcoumarin as a substrate. C. CYP3A4 enzymatic activity was inhibited by 83% with a CYP3A4-specific inhibitor (10 μM ketoconazole). D. Quantitative RT-PCR analysis of the ALBUMIN (ALB) gene on HepaMN cells with exposure to 20 μM rifampicin. E. Quantitative RT-PCR analysis of the α-FETOPROTEIN (AFP) gene on HepaMN cells with exposure to 20 μM rifampicin.

## Discussion

To date, HepaRG is the best hepatocyte line for drug interaction studies and hepatitis virus infection. We herewith introduce a novel hepatocyte line, HepaMN, that was established from a patient with biliary atresia, a childhood disease of the liver in which bile ducts are abnormally narrow or absent. While HepaRG and HepG2 were generated from hepatoma or hepatocellular carcinoma, HepaMN was derived from untransformed hepatocytes, probably hepatic progenitors, like other epithelial progenitors. From this viewpoint, HepaMN cells are the first immortalized hepatocyte cell line with normal function and diploid chromosomes. It is most noteworthy that HepaMN cells exhibited comparable expression levels of the albumin gene with HepaRG, and generated liver-like morphology in vivo. Liver, a multifunctional organ, consists of hepatocytes, pericytes, and stellate cells. Among these cells, hepatocytes play a key role in metabolism and detoxification. However, freshly isolated hepatocytes show considerable variation from lot to lot and are difficult to propagate in vitro. With comparable physiological functions to freshly isolated hepatocytes, the cloned cell line HepaMN was selected from among 11 immortalized hepatocyte clones from Japanese samples (Supplemental Table 3, 4).

HepaMN cells may serve as a tool to analyze in vitro pharmacokinetics and for pharmacology studies of absorption, distribution, metabolism, and excretion; a material to analyze in vitro hepatotoxicity testing with reactive metabolites; a substrate for manufacture of vaccines; and a target cell for infection and replication of hepatitis virus. In the general scheme of model-based prediction from the Food and Drug Administration (FDA) draft guidance, an in vitro induction system is established in cultured human hepatocytes from 3 donors (21). Despite the variability seen among hepatocyte lots, freshly isolated human hepatocytes have been the system of choice for evaluating enzyme induction in vitro, HepaMN cells may also be useful for an in vitro induction system. At the early stages of drug screening, HepaMN cells may have an advantage over HepaRG and Hep G2 and the other systems currently in use, such as freshly isolated hepatocytes, liver microsomes, and cytosolic fractions. Most importantly, HepaMN cells resembled normal hepatocytes with regard to two important hepatic functional features: (i) maintenance of efficient proliferation accompanied by constitutive expression of albumin, and (ii) the ability to display metabolic functions. In addition to the vitro benefits, immunodeficient animals with human HepaMN cells in livers can also be prepared for in vivo hepatotoxicity tests, albeit the low implantation rate of HepaMN cells into the mouse liver through the spleen. In order to generate mice with humanized livers, more efficient implantation methods and routes need to be determined, i.e. cell injection through the portal vein.

## Conclusions

Novel regenerative medicine products such as decellularized liver, liver assistance devices, and bioartificial livers have recently been developed. For pharmaceutical evaluation of these systems, HepaMN cells may be valuable as a cost-effective and relatively easy-to-use substitute for hepatocytes dissociated from livers or from primary cultivation. Most importantly, HepaMN cells possess the outstanding characteristics of immortality and retention of hepatic features even after a long cultivation period.

## Declarations

### Ethics approval and consent to participate

Human cells in this study were performed in full compliance with the Ethical Guidelines for Clinical Studies (Ministry of Health, Labor, and Welfare, Japan). The cells were banked after approval of the Institutional Review Board at the National Institute of Biomedical Innovation (May 9, 2006). Animal experiments were performed according to protocols approved by the Institutional Animal Care and Use Committee of the National Research Institute for Child Health and Development.

### Consent for publication

Not applicable.

### Availability of data and material

The datasets used and/or analysed during the current study are available from the corresponding author on reasonable request.

### Authors’ contributions

AU designed experiments. MN, MT, YO, SI, SH, HM, SO, TO, SE, and TKiy performed experiments. AU, MN, MT, YO, HM, TKiy, SO, TO, and SE analyzed data. AN, MK, and TKiy contributed reagents, materials and analysis tools. AU, MT, MN, HA, TKiy, and TKim discussed the data and manuscript. AU, MN, and TKiy wrote this manuscript.

## Acknowledgements

We would like to express our sincere thanks to M. Ichinose and H. Abe for providing expert technical assistance, to Dr. C. Ketcham for English editing and proofreading, and to E. Suzuki, and K. Saito for secretarial work.

## List of Abbreviations

Not applicable.

## List of Supplemental Information

**Supplemental Figure 1. Implantation of non-immortalized hepatocytes (Hep2001) into NOD/SCID IL-2 receptor** *γ* **-/-(NOG) immunodeficient mice**

Hepatocytes (1.5 × 10^7^ cells) were injected into the thigh muscles of NOG mice. The samples were resected at 4 weeks after the implantation and analyzed for histology and immunohistology. Hepatocytes are shown by dashed lines. A. Hematoxylin-eosin (HE) stain. B to D. Immunohistochemistry of the implanted hepatocytes by using the antibodies to vimentin (B), albumin (C: ALB), and cytokeratin 8/18 (D: CK8/18). The hepatocytes are negative for vimentin and positive for albumin and cytokeratin 8/18. The vimentin-positive, albumin-negative, cytokeratin 8/18-negative cells around vimentin-negative hepatocytes are mesenchymal cells in the sample Hep2001.

**Supplemental Figure 2. Gene induction test for CYP1A1 and CYP1A2 in HepaMN cells** Quantitative RT-PCR analysis of the genes for CYP1A1 and CYP1A2 was performed on HepaMN cells with exposure to omeprazole or β-naphthoflavone at the indicated concentration for 24 h.

**Supplemental Table 1. Primer pairs and experimental conditions for RT-PCR**

**Supplemental Table 2. Liver-associated genes**

**Supplemental Table 3. List of immortalized hepatocytes**

**Supplemental Table 4. List of liver samples**

**Supplemental Table 5. Hepatic differentiation stage of hESCs used for principal component analysis (PCA)**

Human embryonic stem cell (hESC) lines SEES1, SEES4, and SEES5 were stably maintained in XF hESC culture medium containing 85% Knockout DMEM, 15% Knockout Serum Replacement XF CTS, 2 mM GlutaMAX-I, 0.1 mM NEAA, Pen-Strep, 50 µg/mL L-ascorbic acid 2-phosphate, 10 ng/mL heregulin-1β (recombinant human NRG-beta 1/HRG-beta 1 EGF domain), 200 ng/mL recombinant human IGF-1 (LONG R3-IGF-1; Sigma-Aldrich), and 20 ng/mL human bFGF (Akutsu et al., Regenerative Therapy; JCI insight). Undifferentiated hESCs were dissociated using dispase and plated on a dish coated with 0.1% human recombinant type I collagen peptide in 90 mm culture dishes. For hepatic differentiation, hESCs were cultured in XF hESC medium without growth factors (XF-KSR(-)) for 1 day and then in XF-KSR medium, which was replaced after 3 days with the XF hesc medium used as the differentiation medium. The differentiation medium was gently changed every 3–4 days until the indicated day.

