## Supplementary figures and images for "Immortalization of human zone I hepatocytes from biliary atresia with CDK4^R24C^, cyclin D1, and TERT for cytochrome P450 induction testing"

### Supplemental Figure 1

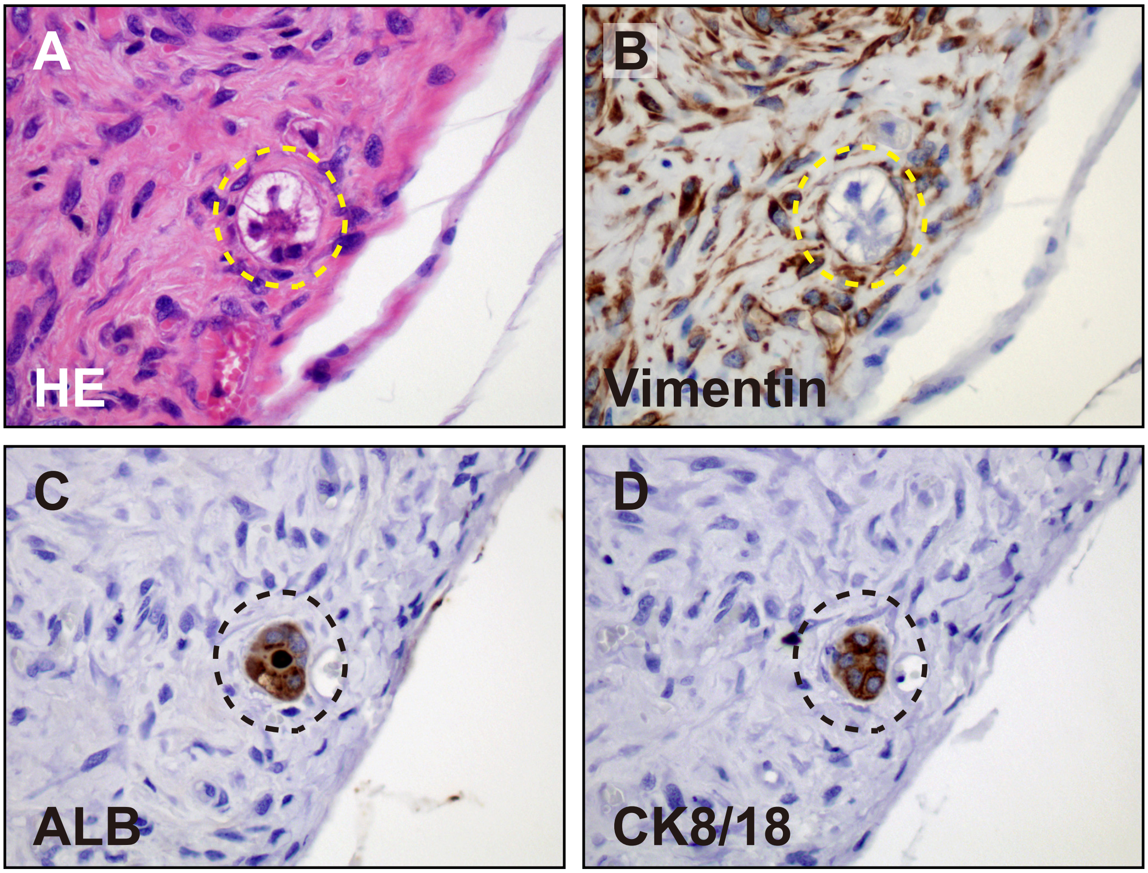


Supplemental Figure 1

### Supplemental Figure 2

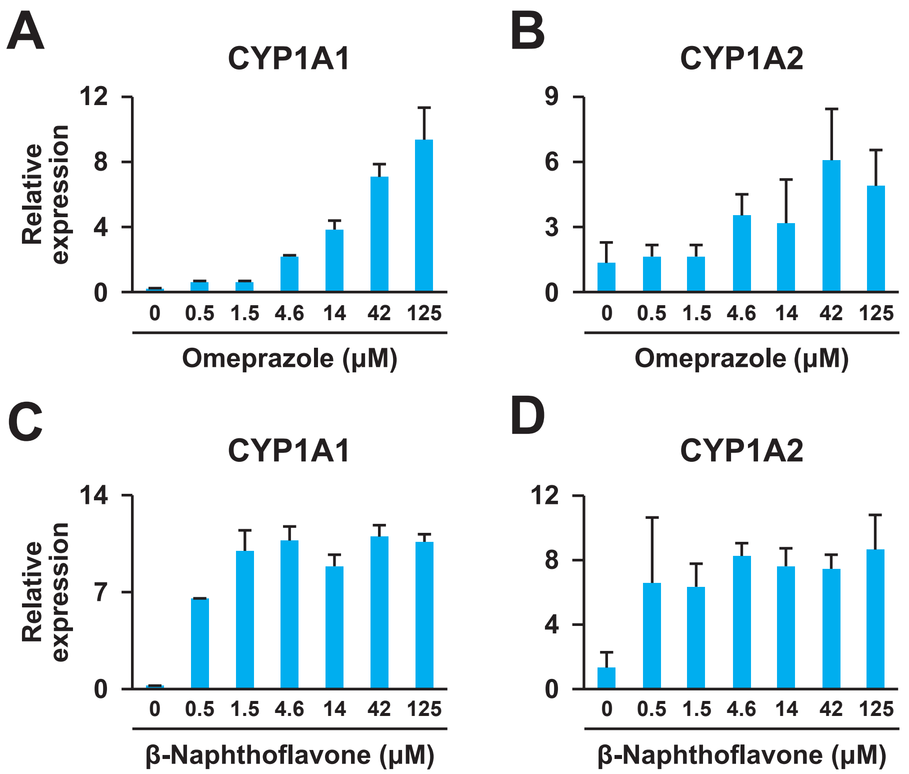


**Supplemental Figure 2**
