## Supplemental Table 1 for "Immortalization of human zone I hepatocytes from biliary atresia with CDK4^R24C^, cyclin D1, and TERT for cytochrome P450 induction testing"

**Supplemental Table 1. Primer pairs and experimental conditions for RT-PCR**

| Gene product | Forward and reverse primers (5'–3') | Expected product size |
|---|---|---|
| AFP | AGCTTGGTGGTGGATGAAAC | 248 |
| | CCCTCTTCAGCAAAGCAGAC | |
| ALB | TGGCACAATGAAGTGGGTAA | 166 |
| | CTGAGCAAAGGCAATCAACA | |
| CYP1A2 | CAATCAGGTGGTGGTGTCAG | 245 |
| | GCTCCTGGACTGTTTTCTGC | |
| CYP2B6 | TCCTTTCTGAGGTTCCGAGA | 416 |
| | TCCCGAAGTCCCTCATAGTG | |
| CYP3A4 | CAAGACCCCTTTGTGGAAAA | 187 |
| | CGAGGCGACTTTCTTTCATC | |
| AAT | GGGAAACTACAGCACCTGGA | 175 |
| | CCCCATTGCTGAAGACCTTA | |
| TDO2 | GGGAACTACCTGCATTTGGA | 222 |
| | GTGCATCCGAGAAACAACCT | |
| OTC | ACCTTCAGGCAGCTACTCCA | 192 |
| | GCCGCTTTTTCTTCTCCTCT | |
| ARG | GGCTGGTCTGCTTGAGAAAC | 240 |
| | ATTGCCAAACTGTGGTCTCC | |
| UBIQUITIN | GGAGCCGAGTGACACCATTG | 346 |
| | CAGGGTACGACCATCTTCCAG | |
