## Supplemental Table 2 for "Immortalization of human zone I hepatocytes from biliary atresia with CDK4^R24C^, cyclin D1, and TERT for cytochrome P450 induction testing"

**Supplemental Table 2. Liver-associated genes**

**A. Developmental markers**

| ProbeName | GeneSymbol | GeneName | Description |
|---|---|---|---|
| A_23_P204395 | AACS | acetoacetyl-CoA synthetase | acetoacetyl-CoA synthetase (AACS) |
| A_33_P3403392 | AACS | acetoacetyl-CoA synthetase | acetoacetyl-CoA synthetase (AACS) |
| A_23_P24515 | ACAT1 | acetyl-CoA acetyltransferase 1 | acetyl-CoA acetyltransferase 1 (ACAT1) |
| A_24_P203678 | ACAT1 | acetyl-CoA acetyltransferase 1 | acetyl-CoA acetyltransferase 1 (ACAT1) |
| A_33_P3253975 | ACAT1 | acetyl-CoA acetyltransferase 1 | acetyl-CoA acetyltransferase 1 (ACAT1) |
| A_23_P200404 | AK2 | adenylate kinase 2 | adenylate kinase 2 (AK2) |
| A_24_P179903 | AK2 | adenylate kinase 2 | adenylate kinase 2 (AK2) |
| A_24_P500891 | AK2 | adenylate kinase 2 | adenylate kinase 2 (AK2) |
| A_33_P3392580 | AK2 | adenylate kinase 2 | adenylate kinase 2 (AK2) |
| A_24_P73577 | ALDH1A2 | aldehyde dehydrogenase 1 family, member A2 | aldehyde dehydrogenase 1 family, member A2 (ALDH1A2) |
| A_32_P18440 | ARID5B | AT rich interactive domain 5B (MRF1-like) | AT rich interactive domain 5B (MRF1-like) (ARID5B) |
| A_33_P3324980 | ARID5B | AT rich interactive domain 5B (MRF1-like) | AT rich interactive domain 5B (MRF1-like) (ARID5B) |
| A_23_P145694 | ASNS | asparagine synthetase (glutamine-hydrolyzing) | asparagine synthetase (glutamine-hydrolyzing) (ASNS) |
| A_23_P31921 | ASS1 | argininosuccinate synthase 1 | argininosuccinate synthase 1 (ASS1) |
| A_33_P3234580 | ASS1 | argininosuccinate synthase 1 | argininosuccinate synthase 1 (ASS1) |
| A_23_P143987 | ATG7 | autophagy related 7 | autophagy related 7 (ATG7) |
| A_24_P944827 | ATG7 | autophagy related 7 | autophagy related 7 (ATG7) |
| A_33_P3252359 | BDH1 | 3-hydroxybutyrate dehydrogenase, type 1 | 3-hydroxybutyrate dehydrogenase, type 1 (BDH1) |
| A_33_P3211138 | CADM1 | cell adhesion molecule 1 | cell adhesion molecule 1 (CADM1) |
| A_33_P3421913 | CADM1 | cell adhesion molecule 1 | cell adhesion molecule 1 (CADM1) |
| A_33_P3745146 | CADM1 | cell adhesion molecule 1 | cell adhesion molecule 1 (CADM1) |
| A_32_P404549 | CCDC39 | coiled-coil domain containing 39 | coiled-coil domain containing 39 (CCDC39) |
| A_24_P160680 | CCDC40 | coiled-coil domain containing 40 | coiled-coil domain containing 40 (CCDC40) |
| A_33_P3302518 | CCDC40 | coiled-coil domain containing 40 | coiled-coil domain containing 40 (CCDC40) |
| A_33_P3253807 | CEBPG | CCAAT/enhancer binding protein (C/EBP), gamma | CCAAT/enhancer binding protein (C/EBP), gamma (CEBPG) |
| A_21_P0011496 | CES1 | carboxylesterase 1 | carboxylesterase 1 (CES1) |
| A_23_P206733 | CES1 | carboxylesterase 1 | carboxylesterase 1 (CES1) |
| A_33_P3241269 | CES1 | carboxylesterase 1 | carboxylesterase 1 (CES1) |
| A_33_P3389704 | CES1 | carboxylesterase 1 | carboxylesterase 1 (CES1) |
| A_23_P214969 | CITED2 | Cbp/p300-interacting transactivator, with Glu/Asp-rich carboxy-terminal doma | Cbp/p300-interacting transactivator, with Glu/Asp-rich carboxy-terminal domain, 2 (CITED2) |
| A_33_P3213374 | CITED2 | Cbp/p300-interacting transactivator, with Glu/Asp-rich carboxy-terminal doma | Cbp/p300-interacting transactivator, with Glu/Asp-rich carboxy-terminal domain, 2 (CITED2) |
| A_33_P3327673 | COBL | cordon-bleu WH2 repeat protein | cordon-bleu WH2 repeat protein (COBL) |
| A_33_P3725227 | COBL | cordon-bleu WH2 repeat protein | cordon-bleu WH2 repeat protein (COBL) |
| A_33_P3242798 | CPS1 | carbamoyl-phosphate synthase 1, mitochondrial | carbamoyl-phosphate synthase 1, mitochondrial (CPS1) |
| A_23_P29495 | CTNNB1 | catenin (cadherin-associated protein), beta 1, 88kDa | catenin (cadherin-associated protein), beta 1, 88kDa (CTNNB1) |
| A_33_P3421695 | CTNNB1 | catenin (cadherin-associated protein), beta 1, 88kDa | catenin (cadherin-associated protein), beta 1, 88kDa (CTNNB1) |
| A_23_P163402 | CYP1A1 | cytochrome P450, family 1, subfamily A, polypeptide 1 | cytochrome P450, family 1, subfamily A, polypeptide 1 (CYP1A1) |
| A_23_P130753 | DBP | D site of albumin promoter (albumin D-box) binding protein | D site of albumin promoter (albumin D-box) binding protein (DBP) |
| A_23_P54612 | DNAAF1 | dynein, axonemal, assembly factor 1 | dynein, axonemal, assembly factor 1 (DNAAF1) |

| | | | |
|---|---|---|---|
| A_33_P3368388 | DNAAF1 | dynein, axonemal, assembly factor 1 | dynein, axonemal, assembly factor 1 (DNAAF1) |
| A_32_P210202 | E2F7 | E2F transcription factor 7 | E2F transcription factor 7 (E2F7) |
| A_33_P3318661 | E2F7 | E2F transcription factor 7 | E2F transcription factor 7 (E2F7) |
| A_23_P35871 | E2F8 | E2F transcription factor 8 | E2F transcription factor 8 (E2F8) |
| A_23_P215790 | EGFR | epidermal growth factor receptor | epidermal growth factor receptor (EGFR) |
| A_33_P3351944 | EGFR | epidermal growth factor receptor | epidermal growth factor receptor (EGFR) |
| A_33_P3351955 | EGFR | epidermal growth factor receptor | epidermal growth factor receptor (EGFR) |
| A_24_P397150 | GAK | cyclin G associated kinase | cyclin G associated kinase (GAK) |
| A_33_P3254013 | GAK | cyclin G associated kinase | cyclin G associated kinase (GAK) |
| A_23_P429184 | GNPNAT1 | glucosamine-phosphate N-acetyltransferase 1 | glucosamine-phosphate N-acetyltransferase 1 (GNPNAT1) |
| A_33_P3276713 | HGF | hepatocyte growth factor (hepapoietin A; scatter factor) | hepatocyte growth factor (hepapoietin A; scatter factor) (HGF) |
| A_33_P3276718 | HGF | hepatocyte growth factor (hepapoietin A; scatter factor) | hepatocyte growth factor (hepapoietin A; scatter factor) (HGF) |
| A_23_P47034 | HHEX | hematopoietically expressed homeobox | hematopoietically expressed homeobox (HHEX) |
| A_23_P126266 | HLX | H2.0-like homeobox | H2.0-like homeobox (HLX) |
| A_24_P63522 | HMGCS1 | 3-hydroxy-3-methylglutaryl-CoA synthase 1 (soluble) | 3-hydroxy-3-methylglutaryl-CoA synthase 1 (soluble) (HMGCS1) |
| A_33_P3299119 | HNF1A | HNF1 homeobox A | HNF1 homeobox A (HNF1A) |
| A_33_P3299122 | HNF1A | HNF1 homeobox A | HNF1 homeobox A (HNF1A) |
| A_33_P3328312 | HNF1A | HNF1 homeobox A | HNF1 homeobox A (HNF1A) |
| A_23_P406782 | HPN | hepsin | hepsin (HPN) |
| A_24_P128524 | ICMT | isoprenylcysteine carboxyl methyltransferase | isoprenylcysteine carboxyl methyltransferase (ICMT) |
| A_33_P3381022 | ICMT | isoprenylcysteine carboxyl methyltransferase | isoprenylcysteine carboxyl methyltransferase (ICMT) |
| A_23_P48339 | IFT88 | intraflagellar transport 88 | intraflagellar transport 88 (IFT88) |
| A_32_P178800 | ITGA2 | integrin, alpha 2 (CD49B, alpha 2 subunit of VLA-2 receptor) | integrin, alpha 2 (CD49B, alpha 2 subunit of VLA-2 receptor) (ITGA2) |
| A_23_P142389 | LSR | lipolysis stimulated lipoprotein receptor | lipolysis stimulated lipoprotein receptor (LSR) |
| A_33_P3214466 | MESP1 | mesoderm posterior basic helix-loop-helix transcription factor 1 | mesoderm posterior basic helix-loop-helix transcription factor 1 (MESP1) |
| A_23_P54556 | MKL2 | MKL/myocardin-like 2 | MKL/myocardin-like 2 (MKL2) |
| A_24_P375205 | MKL2 | MKL/myocardin-like 2 | cDNA FLJ36258 fis, clone THYMU2002450 |
| A_21_P0012236 | NF1 | neurofibromin 1 | neurofibromin 1 (NF1) |
| A_24_P1919 | NF1 | neurofibromin 1 | neurofibromin 1 (NF1) |
| A_24_P917026 | NF1 | neurofibromin 1 | neurofibromin 1 (NF1) |
| A_33_P3240348 | NF1 | neurofibromin 1 | neurofibromin 1 (NF1) |
| A_33_P3381490 | NKX2-8 | NK2 homeobox 8 | NK2 homeobox 8 (NKX2-8) |
| A_33_P3228072 | NPHP3 | nephronophthisis 3 (adolescent) | nephronophthisis 3 (adolescent) (NPHP3) |
| A_33_P3228102 | NPHP3 | nephronophthisis 3 (adolescent) | nephronophthisis 3 (adolescent) (NPHP3) |
| A_32_P142440 | PCSK9 | proprotein convertase subtilisin/kexin type 9 | proprotein convertase subtilisin/kexin type 9 (PCSK9) |
| A_23_P502371 | PHF2 | PHD finger protein 2 | PHD finger protein 2 (PHF2) |
| A_24_P106112 | PKD2 | polycystic kidney disease 2 (autosomal dominant) | polycystic kidney disease 2 (autosomal dominant) (PKD2) |
| A_24_P396197 | PRKCSH | protein kinase C substrate 80K-H | protein kinase C substrate 80K-H (PRKCSH) |
| A_24_P88266 | PROX1 | prospero homeobox 1 | prospero homeobox 1 (PROX1) |
| A_24_P191847 | PTCD2 | pentatricopeptide repeat domain 2 | pentatricopeptide repeat domain 2 (PTCD2) |
| A_33_P3371819 | PTCD2 | pentatricopeptide repeat domain 2 | pentatricopeptide repeat domain 2 (PTCD2) |
| A_23_P207842 | RARA | retinoic acid receptor, alpha | retinoic acid receptor, alpha (RARA) |

| | | | |
|---|---|---|---|
| A_32_P5251 | RARA | retinoic acid receptor, alpha | retinoic acid receptor, alpha (RARA) |
| A_23_P9056 | RB1CC1 | RB1-inducible coiled-coil 1 | RB1-inducible coiled-coil 1 (RB1CC1) |
| A_23_P104689 | RELA | v-rel avian reticuloendotheliosis viral oncogene homolog A | v-rel avian reticuloendotheliosis viral oncogene homolog A (RELA) |
| A_33_P3209433 | RELA | v-rel avian reticuloendotheliosis viral oncogene homolog A | v-rel avian reticuloendotheliosis viral oncogene homolog A (RELA) |
| A_33_P3274069 | RHBDD3 | rhomboid domain containing 3 | rhomboid domain containing 3 (RHBDD3) |
| A_33_P3330731 | RHBDD3 | rhomboid domain containing 3 | rhomboid domain containing 3 (RHBDD3) |
| A_33_P3302861 | RPGRIP1L | RPGRIP1-like | RPGRIP1-like (RPGRIP1L) |
| A_24_P34155 | RUNX1 | runt-related transcription factor 1 | runt-related transcription factor 1 (RUNX1) |
| A_24_P96403 | RUNX1 | runt-related transcription factor 1 | runt-related transcription factor 1 (RUNX1) |
| A_33_P3211804 | RUNX1 | runt-related transcription factor 1 | runt-related transcription factor 1 (RUNX1) |
| A_33_P3211809 | RUNX1 | runt-related transcription factor 1 | runt-related transcription factor 1 (RUNX1) |
| A_33_P3211818 | RUNX1 | runt-related transcription factor 1 | runt-related transcription factor 1 (RUNX1) |
| A_23_P214977 | SEC63 | SEC63 homolog (S. cerevisiae) | SEC63 homolog (S. cerevisiae) (SEC63) |
| A_24_P123720 | SEC63 | SEC63 homolog (S. cerevisiae) | SEC63 homolog (S. cerevisiae) (SEC63) |
| A_23_P150768 | SLCO2B1 | solute carrier organic anion transporter family, member 2B1 | solute carrier organic anion transporter family, member 2B1 (SLCO2B1) |
| A_23_P48936 | SMAD3 | SMAD family member 3 | SMAD family member 3 (SMAD3) |
| A_23_P134176 | SOD2 | superoxide dismutase 2, mitochondrial | superoxide dismutase 2, mitochondrial (SOD2) |
| A_33_P3380867 | STAT5B | signal transducer and activator of transcription 5B | signal transducer and activator of transcription 5B (STAT5B) |
| A_23_P200780 | TGFBR3 | transforming growth factor, beta receptor III | transforming growth factor, beta receptor III (TGFBR3) |
| A_24_P157926 | TNFAIP3 | tumor necrosis factor, alpha-induced protein 3 | tumor necrosis factor, alpha-induced protein 3 (TNFAIP3) |
| A_23_P47704 | UCP2 | uncoupling protein 2 (mitochondrial, proton carrier) | uncoupling protein 2 (mitochondrial, proton carrier) (UCP2) |
| A_23_P327910 | ZIC3 | Zic family member 3 | Zic family member 3 (ZIC3) |
| A_33_P3250861 | ZIC3 | Zic family member 3 | Zic family member 3 (ZIC3) |

**B. Mature hepatocyte markers**

| ProbeName | GeneSymbol | GeneName | Description |
|---|---|---|---|
| A_23_P257834 | ALB | albumin | albumin (ALB) |
| A_33_P3242798 | CPS1 | carbamoyl-phosphate synthase 1, mitochondrial | carbamoyl-phosphate synthase 1, mitochondrial (CPS1) |
| A_32_P73821 | CSDE1 | cold shock domain containing E1, RNA-binding | cold shock domain containing E1, RNA-binding (CSDE1) |
| A_23_P129169 | CYP11A1 | cytochrome P450, family 11, subfamily A, polypeptide 1 | cytochrome P450, family 11, subfamily A, polypeptide 1 (CYP11A1) |
| A_24_P329424 | CYP11B1 | cytochrome P450, family 11, subfamily B, polypeptide 1 | cytochrome P450, family 11, subfamily B, polypeptide 1 (CYP11B1) |
| A_23_P215997 | CYP11B2 | cytochrome P450, family 11, subfamily B, polypeptide 2 | cytochrome P450, family 11, subfamily B, polypeptide 2 (CYP11B2) |
| A_33_P3376478 | CYP17A1 | cytochrome P450, family 17, subfamily A, polypeptide 1 | cytochrome P450, family 17, subfamily A, polypeptide 1 (CYP17A1) |
| A_24_P920646 | CYP19A1 | cytochrome P450, family 19, subfamily A, polypeptide 1 | cytochrome P450, family 19, subfamily A, polypeptide 1 (CYP19A1) |
| A_32_P86289 | CYP19A1 | cytochrome P450, family 19, subfamily A, polypeptide 1 | cytochrome P450, family 19, subfamily A, polypeptide 1 (CYP19A1) |
| A_23_P37410 | CYP19A1 | cytochrome P450, family 19, subfamily A, polypeptide 1 | cytochrome P450, family 19, subfamily A, polypeptide 1 (CYP19A1) |
| A_33_P3351371 | CYP19A1 | cytochrome P450, family 19, subfamily A, polypeptide 1 | cytochrome P450, family 19, subfamily A, polypeptide 1 (CYP19A1) |
| A_23_P163402 | CYP1A1 | cytochrome P450, family 1, subfamily A, polypeptide 1 | cytochrome P450, family 1, subfamily A, polypeptide 1 (CYP1A1) |
| A_33_P3253747 | CYP1A2 | cytochrome P450, family 1, subfamily A, polypeptide 2 | cytochrome P450, family 1, subfamily A, polypeptide 2 (CYP1A2) |
| A_23_P209625 | CYP1B1 | cytochrome P450, family 1, subfamily B, polypeptide 1 | cytochrome P450, family 1, subfamily B, polypeptide 1 (CYP1B1) |
| A_33_P3290343 | CYP1B1 | cytochrome P450, family 1, subfamily B, polypeptide 1 | cytochrome P450, family 1, subfamily B, polypeptide 1 (CYP1B1) |
| A_22_P00006287 | CYP1B1-AS1 | CYP1B1 antisense RNA 1 | CYP1B1 antisense RNA 1 (CYP1B1-AS1) |
| A_19_P00807643 | CYP1B1-AS1 | CYP1B1 antisense RNA 1 | CYP1B1 antisense RNA 1 (CYP1B1-AS1) |

| | | | |
|---|---|---|---|
| A_23_P56894 | CYP20A1 | cytochrome P450, family 20, subfamily A, polypeptide 1 | cytochrome P450, family 20, subfamily A, polypeptide 1 (CYP20A1) |
| A_33_P3371175 | CYP20A1 | cytochrome P450, family 20, subfamily A, polypeptide 1 | cytochrome P450, family 20, subfamily A, polypeptide 1 (CYP20A1) |
| A_23_P257478 | CYP21A2 | cytochrome P450, family 21, subfamily A, polypeptide 2 | cytochrome P450, family 21, subfamily A, polypeptide 2 (CYP21A2) |
| A_33_P3411279 | CYP21A2 | cytochrome P450, family 21, subfamily A, polypeptide 2 | cytochrome P450, family 21, subfamily A, polypeptide 2 (CYP21A2) |
| A_23_P28815 | CYP24A1 | cytochrome P450, family 24, subfamily A, polypeptide 1 | cytochrome P450, family 24, subfamily A, polypeptide 1 (CYP24A1) |
| A_33_P3369401 | CYP24A1 | cytochrome P450, family 24, subfamily A, polypeptide 1 | cytochrome P450, family 24, subfamily A, polypeptide 1 (CYP24A1) |
| A_23_P138655 | CYP26A1 | cytochrome P450, family 26, subfamily A, polypeptide 1 | cytochrome P450, family 26, subfamily A, polypeptide 1 (CYP26A1) |
| A_23_P210109 | CYP26B1 | cytochrome P450, family 26, subfamily B, polypeptide 1 | cytochrome P450, family 26, subfamily B, polypeptide 1 (CYP26B1) |
| A_33_P3361422 | CYP27A1 | cytochrome P450, family 27, subfamily A, polypeptide 1 | cytochrome P450, family 27, subfamily A, polypeptide 1 (CYP27A1) |
| A_23_P36397 | CYP27B1 | cytochrome P450, family 27, subfamily B, polypeptide 1 | cytochrome P450, family 27, subfamily B, polypeptide 1 (CYP27B1) |
| A_23_P55779 | CYP2A13 | cytochrome P450, family 2, subfamily A, polypeptide 13 | cytochrome P450, family 2, subfamily A, polypeptide 13 (CYP2A13) |
| A_23_P27528 | CYP2A7 | cytochrome P450, family 2, subfamily A, polypeptide 7 | cytochrome P450, family 2, subfamily A, polypeptide 7 (CYP2A7) |
| A_24_P339514 | CYP2B6 | cytochrome P450, family 2, subfamily B, polypeptide 6 | cytochrome P450, family 2, subfamily B, polypeptide 6 (CYP2B6) |
| A_23_P208373 | CYP2B6 | cytochrome P450, family 2, subfamily B, polypeptide 6 | cytochrome P450, family 2, subfamily B, polypeptide 6 (CYP2B6) |
| A_23_P52480 | CYP2C18 | cytochrome P450, family 2, subfamily C, polypeptide 18 | cytochrome P450, family 2, subfamily C, polypeptide 18 (CYP2C18) |
| A_33_P3326075 | CYP2C19 | cytochrome P450, family 2, subfamily C, polypeptide 19 | cytochrome P450, family 2, subfamily C, polypeptide 19 (CYP2C19) |
| A_23_P161368 | CYP2C8 | cytochrome P450, family 2, subfamily C, polypeptide 8 | cytochrome P450, family 2, subfamily C, polypeptide 8 (CYP2C8) |
| A_23_P12767 | CYP2C9 | cytochrome P450, family 2, subfamily C, polypeptide 9 | cytochrome P450, family 2, subfamily C, polypeptide 9 (CYP2C9) |
| A_23_P155123 | CYP2D6 | cytochrome P450, family 2, subfamily D, polypeptide 6 | cytochrome P450, family 2, subfamily D, polypeptide 6 (CYP2D6) |
| A_23_P143734 | CYP2D6 | cytochrome P450, family 2, subfamily D, polypeptide 6 | cytochrome P450, family 2, subfamily D, polypeptide 6 (CYP2D6) |
| A_24_P394940 | CYP2E1 | cytochrome P450, family 2, subfamily E, polypeptide 1 | cytochrome P450, family 2, subfamily E, polypeptide 1 (CYP2E1) |
| A_23_P89981 | CYP2F1 | cytochrome P450, family 2, subfamily F, polypeptide 1 | cytochrome P450, family 2, subfamily F, polypeptide 1 (CYP2F1) |
| A_23_P103486 | CYP2J2 | cytochrome P450, family 2, subfamily J, polypeptide 2 | cytochrome P450, family 2, subfamily J, polypeptide 2 (CYP2J2) |
| A_23_P202860 | CYP2R1 | cytochrome P450, family 2, subfamily R, polypeptide 1 | cytochrome P450, family 2, subfamily R, polypeptide 1 (CYP2R1) |
| A_21_P0014273 | CYP2R1 | cytochrome P450, family 2, subfamily R, polypeptide 1 | cytochrome P450, family 2, subfamily R, polypeptide 1 (CYP2R1) |
| A_23_P101374 | CYP2S1 | cytochrome P450, family 2, subfamily S, polypeptide 1 | cytochrome P450, family 2, subfamily S, polypeptide 1 |
| A_33_P3348782 | CYP2S1 | cytochrome P450, family 2, subfamily S, polypeptide 1 | cytochrome P450, family 2, subfamily S, polypeptide 1 |
| A_33_P3252605 | CYP2U1 | cytochrome P450, family 2, subfamily U, polypeptide 1 | cytochrome P450, family 2, subfamily U, polypeptide 1 (CYP2U1) |
| A_33_P3252612 | CYP2W1 | cytochrome P450, family 2, subfamily W, polypeptide 1 | cytochrome P450, family 2, subfamily W, polypeptide 1 (CYP2W1) |
| A_23_P133712 | CYP39A1 | cytochrome P450, family 39, subfamily A, polypeptide 1 | cytochrome P450, family 39, subfamily A, polypeptide 1 (CYP39A1) |
| A_33_P3251342 | CYP3A4 | cytochrome P450, family 3, subfamily A, polypeptide 4 | cytochrome P450, family 3, subfamily A, polypeptide 4 (CYP3A4) |
| A_23_P215828 | CYP3A43 | cytochrome P450, family 3, subfamily A, polypeptide 43 | cytochrome P450, family 3, subfamily A, polypeptide 43 (CYP3A43) |
| A_23_P8801 | CYP3A5 | cytochrome P450, family 3, subfamily A, polypeptide 5 | cytochrome P450, family 3, subfamily A, polypeptide 5 (CYP3A5) |
| A_33_P3249746 | CYP3A5 | cytochrome P450, family 3, subfamily A, polypeptide 5 | cytochrome P450, family 3, subfamily A, polypeptide 5 (CYP3A5) |
| A_23_P358917 | CYP3A7 | cytochrome P450, family 3, subfamily A, polypeptide 7 | cytochrome P450, family 3, subfamily A, polypeptide 7 (CYP3A7) |
| A_33_P3318117 | CYP3A7 | cytochrome P450, family 3, subfamily A, polypeptide 7 | cytochrome P450, family 3, subfamily A, polypeptide 7 (CYP3A7) |
| A_23_P48784 | CYP46A1 | cytochrome P450, family 46, subfamily A, polypeptide 1 | cytochrome P450, family 46, subfamily A, polypeptide 1 |
| A_33_P3337604 | CYP46A1 | cytochrome P450, family 46, subfamily A, polypeptide 1 | cytochrome P450, family 46, subfamily A, polypeptide 1 |
| A_24_P191013 | CYP4A11 | cytochrome P450, family 4, subfamily A, polypeptide 11 | cytochrome P450, family 4, subfamily F, polypeptide 11 (CYP4F11) |
| A_33_P3303474 | CYP4A11 | cytochrome P450, family 4, subfamily A, polypeptide 11 | cytochrome P450, family 4, subfamily F, polypeptide 11 (CYP4F11) |
| A_23_P114713 | CYP4B1 | cytochrome P450, family 4, subfamily B, polypeptide 1 | cytochrome P450, family 4, subfamily B, polypeptide 1 (CYP4B1) |
| A_21_P0000967 | CYP4B1 | cytochrome P450, family 4, subfamily B, polypeptide 1 | cytochrome P450, family 4, subfamily B, polypeptide 1 |

| | | | |
|---|---|---|---|
| A_24_P42693 | CYP4F11 | cytochrome P450, family 4, subfamily F, polypeptide 11 | cytochrome P450, family 4, subfamily F, polypeptide 11 (CYP4F11) |
| A_23_P39315 | CYP4F11 | cytochrome P450, family 4, subfamily F, polypeptide 11 | cytochrome P450, family 4, subfamily F, polypeptide 11 (CYP4F11) |
| A_23_P108280 | CYP4F12 | cytochrome P450, family 4, subfamily F, polypeptide 12 | cytochrome P450, family 4, subfamily F, polypeptide 12 (CYP4F12) |
| A_23_P50710 | CYP4F2 | cytochrome P450, family 4, subfamily F, polypeptide 2 | cytochrome P450, family 4, subfamily F, polypeptide 2 (CYP4F2) |
| A_33_P3359017 | CYP4F2 | cytochrome P450, family 4, subfamily F, polypeptide 2 | cytochrome P450, family 4, subfamily F, polypeptide 2 (CYP4F2) |
| A_24_P331150 | CYP4F22 | cytochrome P450, family 4, subfamily F, polypeptide 22 | cytochrome P450, family 4, subfamily F, polypeptide 22 (CYP4F22) |
| A_33_P3294277 | CYP4F3 | cytochrome P450, family 4, subfamily F, polypeptide 3 | cytochrome P450, family 4, subfamily F, polypeptide 3 (CYP4F3) |
| A_23_P39881 | CYP4F30P | cytochrome P450, family 4, subfamily F, polypeptide 30, pseudogene | cytochrome P450, family 4, subfamily F, polypeptide 30, pseudogene (CYP4F30P) |
| A_24_P28811 | CYP4F62P | cytochrome P450, family 4, subfamily F, polypeptide 62, pseudogene | cytochrome P450, family 4, subfamily F, polypeptide 62, pseudogene (CYP4F62P) |
| A_23_P131060 | CYP4F8 | cytochrome P450, family 4, subfamily F, polypeptide 8 | cytochrome P450, family 4, subfamily F, polypeptide 8 (CYP4F8) |
| A_24_P945228 | CYP4V2 | cytochrome P450, family 4, subfamily V, polypeptide 2 | cytochrome P450, family 4, subfamily V, polypeptide 2 (CYP4V2) |
| A_24_P293530 | CYP4X1 | cytochrome P450, family 4, subfamily X, polypeptide 1 | cytochrome P450, family 4, subfamily X, polypeptide 1 (CYP4X1) |
| A_23_P103971 | CYP4Z1 | cytochrome P450, family 4, subfamily Z, polypeptide 1 | cytochrome P450, family 4, subfamily Z, polypeptide 1 (CYP4Z1) |
| A_33_P3279880 | CYP4Z1 | cytochrome P450, family 4, subfamily Z, polypeptide 1 | cytochrome P450, family 4, subfamily Z, polypeptide 1 (CYP4Z1) |
| A_21_P0010756 | CYP4Z1 | cytochrome P450, family 4, subfamily Z, polypeptide 1 | cytochrome P450, family 4, subfamily Z, polypeptide 1 (CYP4Z1) |
| A_24_P145529 | CYP4Z2P | cytochrome P450, family 4, subfamily Z, polypeptide 2, pseudogene | cDNA FLJ40054 fis, clone TBAES2000315 |
| A_21_P0010757 | CYP4Z2P | cytochrome P450, family 4, subfamily Z, polypeptide 2, pseudogene | cytochrome P450 (CYP4Z2P) |
| A_24_P130041 | CYP51A1 | cytochrome P450, family 51, subfamily A, polypeptide 1 | cytochrome P450, family 51, subfamily A, polypeptide 1 (CYP51A1) |
| A_23_P146198 | CYP7A1 | cytochrome P450, family 7, subfamily A, polypeptide 1 | cytochrome P450, family 7, subfamily A, polypeptide 1 (CYP7A1) |
| A_23_P169092 | CYP7B1 | cytochrome P450, family 7, subfamily B, polypeptide 1 | cytochrome P450, family 7, subfamily B, polypeptide 1 (CYP7B1) |
| A_24_P208704 | CYP8B1 | cytochrome P450, family 8, subfamily B, polypeptide 1 | cytochrome P450, family 8, subfamily B, polypeptide 1 (CYP8B1) |
| A_33_P3733417 | DRD2 | dopamine receptor D2 | dopamine receptor D2 (DRD2) |
| A_32_P210202 | E2F7 | E2F transcription factor 7 | E2F transcription factor 7 (E2F7) |
| A_33_P3318661 | E2F7 | E2F transcription factor 7 | E2F transcription factor 7 (E2F7) |
| A_23_P35871 | E2F8 | E2F transcription factor 8 | E2F transcription factor 8 (E2F8) |
| A_23_P144843 | ESM1 | endothelial cell-specific molecule 1 | endothelial cell-specific molecule 1 (ESM1) |
| A_23_P160742 | GLMN | glomulin, FKBP associated protein | glomulin, FKBP associated protein (GLMN) |
| A_23_P47034 | HHEX | hematopoietically expressed homeobox | hematopoietically expressed homeobox (HHEX) |
| A_23_P406782 | HPN | hepsin | hepsin (HPN) |
| A_32_P178800 | ITGA2 | integrin, alpha 2 (CD49B, alpha 2 subunit of VLA-2 receptor) | integrin, alpha 2 (CD49B, alpha 2 subunit of VLA-2 receptor) (ITGA2) |
| A_33_P3214466 | MESP1 | mesoderm posterior basic helix-loop-helix transcription factor 1 | mesoderm posterior basic helix-loop-helix transcription factor 1 (MESP1) |
| A_23_P58763 | PELO | pelota homolog (Drosophila) | pelota homolog (Drosophila) (PELO) |
| A_24_P88266 | PROX1 | prospero homeobox 1 | prospero homeobox 1 (PROX1) |
| A_24_P157926 | TNFAIP3 | tumor necrosis factor, alpha-induced protein 3 | tumor necrosis factor, alpha-induced protein 3 (TNFAIP3) |
