## Supplemental Table 3 for "Immortalization of human zone I hepatocytes from biliary atresia with CDK4^R24C^, cyclin D1, and TERT for cytochrome P450 induction testing"

**Supplemental Table 3. List of immortalized hepatocytes**

| Sample name | Introduced gene | | | | Details |
|---|---|---|---|---|---|
| | *TERT* | *CDK4R24C* | *Cyclin D1* | *Tet-Off* | |
| Hep2004 | + | + | + | + | *543:CSII-CMV-hTERT(MOI=3),1718:CSII-CMV-Tet-Off Advanced(MOI=3),1728:CSII-TRE-Tight-hCDK4R24C(MOI=3),1727:CSII-TRE-Tight-cyclin D1(MOI=3)* |
| Hep2013 | + | + | + | + | *543:CSII-CMV-hTERT(MOI=3),1718:CSII-CMV-Tet-Off Advanced(MOI=3),1728:CSII-TRE-Tight-hCDK4R24C(MOI=3),1727:CSII-TRE-Tight-cyclin D1(MOI=3)* |
| Hep2017 | + | + | + | - | *543:CSII-CMV-hTERT,1048:CSII-CMV-hCDK4R24C,1172:CSII-CMV-cyclin D1* |
| HepaMN (Hep2018) | + | + | + | + | *543:CSII-CMV-hTERT(MOI=3),1718:CSII-CMV-Tet-Off Advanced(MOI=5),1728:CSII-TRE-Tight-hCDK4R24C(MOI=10),1727:CSII-TRE-Tight-cyclin D1(MOI=10)* |
| Hep2020 | + | + | + | + | *543:CSII-CMV-hTERT(MOI=3),1718:CSII-CMV-Tet-Off Advanced(MOI=5),1728:CSII-TRE-Tight-hCDK4R24C(MOI=10),1727:CSII-TRE-Tight-cyclin D1(MOI=10)* |
| Hep2040 | + | + | + | - | *543:CSII-CMV-hTERT,1048:CSII-CMV-hCDK4R24C,1172:CSII-CMV-cyclin D1* |
| Hep2044 | + | + | + | + | *543:CSII-CMV-hTERT,1718:CSII-CMV-Tet-Off Advanced,1728:CSII-TRE-Tight-hCDK4R24C,1727:CSII-TRE-Tight-cyclin D1* |
| Hep2045 | + | + | + | + | *543:CSII-CMV-hTERT(MOI=3),1718:CSII-CMV-Tet-Off Advanced(MOI=5),1728:CSII-TRE-Tight-hCDK4R24C(MOI=10),1727:CSII-TRE-Tight-cyclin D1(MOI=10)* |
| Hep2022D | + | + | + | + | *543:CSII-CMV-hTERT(MOI=3),1718:CSII-CMV-Tet-Off Advanced(MOI=5),1728:CSII-TRE-Tight-hCDK4R24C(MOI=10),1727:CSII-TRE-Tight-cyclin D1(MOI=10)* |
| Hep2023D | + | + | + | - | *543:CSII-CMV-hTERT,1048:CSII-CMV-hCDK4R24C(MOI>10?),1172:CSII-CMV-cyclin D1(MOI>10?)* |
| Hep2039D | + | + | + | - | *543:CSII-CMV-hTERT(MOI-3),1048:CSII-CMV-hCDK4R24C(MOI=3),1172:CSII-CMV-cyclin D1(MOI=3)* |

The contructions are detailed in Gene Therapy, 18:857, 2011.
