## Supplemental Table 4 for "Immortalization of human zone I hepatocytes from biliary atresia with CDK4^R24C^, cyclin D1, and TERT for cytochrome P450 induction testing"

**Supplemental Table 4. List of liver samples**

| Sample name | Age | Sex | Recipient or Donor | Disease |
|---|---|---|---|---|
| Hep2001 | | | | MMA |
| Hep2002 | | | R | CPS1D |
| Hep2003 | 14 years and 2 months | M | R | CAPV |
| Hep2004 | | | R | BA |
| Hep2005R | 7 months | F | R | BA |
| Hep2005D | 30 years | F | D | Healthy liver |
| Hep2006 | 2 years and 8 months | F | R | BA |
| Hep2007 | 1 year and 1 month | F | R | BA |
| Hep2008 | 9 months | F | R | BA |
| Hep2009R | 5 months | F | R | BA |
| Hep2009D | 31 years | M | D | Healthy liver |
| Hep2010 | | | R | OTCD |
| Hep2011 | 9 months | F | R | Alagile syndrome |
| Hep2012 | 8 months | F | R | BA |
| Hep2013 | 3 years and 10 months | F | R | BA |
| Hep2014 | 2 years and 1 month | F | R | Propionemia |
| Hep2015R | 1 year and 1 month | F | R | BA |
| Hep2015D | 40 years | M | D | Healthy liver |
| Hep2016 | 12 years and 5 months | F | R | BA |
| Hep2017 | 1 year and 3 months | F | R | GSDIb |
| Hep2018 | 4 years | F | R | BA |
| Hep2019 | 9 months | M | R | BA |
| Hep2020 | 6 years | M | R | CHF |
| Hep2021R | 11 years and 5 months | M | R | GSDI |
| Hep2022D | 26 years | M | D | Healthy liver |
| Hep2023D | 32 years | M | D | Healthy liver |
| Hep2023R | 10 months | F | R | BA |
| Hep2024D | 21 years | F | D | Healthy liver |
| Hep2039D | 36 years | F | D | Healthy liver |
| Hep2040 | 7 months | M | R | IHC |
| Hep2044 | 5 months | F | R | BA |
| Hep2045 | 5 years | M | R | BA |

MMA: Methylmalonic acidemia
CPS1D: Carbamyl phosphate synthetase deficiency
CAPV: Congenital absence of the portal vein
BA: Biliary atresia
OTCD: Ornithine transcarbamylase deficiency
GSD: Glycogen storage disease
CHF: Congenital hepatic fibrosis
IHC: Intrahepatic cholestasis
