## Supplemental Table 5 for "Immortalization of human zone I hepatocytes from biliary atresia with CDK4^R24C^, cyclin D1, and TERT for cytochrome P450 induction testing"

**Supplemental Table 5. Hepatic differentiation stage of hESCs used for principal component analysis (PCA)**

| Sample Number | Sample Name | Days cultivated after the start of the hepatic differentiation | Estimated Differentiation Stage | ESCs |
|---|---|---|---|---|
| 1 | SEES1-LGR5 | 0 days | Undifferentiated cells | SEES1 |
| 2 | SEES4 | 0 days | Undifferentiated cells | SEES4 |
| 4 | SEES5-P56 | 50 days | Mature hepatocyte-like cells | SEES5 |
| 5 | LGR5F+I d7 | 7 days | Undifferentiated endodermal cells | SEES1 |
| 6 | LGR5F+I d14 | 14 days | Immature hepatocyte-like cells | SEES1 |
| 7 | LGR5F+H d7 | 7 days | Undifferentiated endodermal cells | SEES1 |
| 8 | LGR5F+H d14 | 14 days | Immature hepatocyte-like cells | SEES1 |
| 9 | LGR5H+I d7 | 7 days | Undifferentiated endodermal cells | SEES1 |
| 10 | LGR5H+I d14 | 7 days | Undifferentiated endodermal cells | SEES1 |
| 11 | LGR5XF- d7 | 7 days | Undifferentiated endodermal cells | SEES1 |
| 12 | LGR5XF- d14 | 14 days | Immature hepatocyte-like cells | SEES1 |
| 13 | LGR5XF32 d30 | 30 days | Mature hepatocyte-like cells | SEES1 |
| 14 | LGR5XF32 d60 | 60 days | Mature hepatocyte-like cells | SEES1 |
| 15 | SEES4-EB | 14 days | Immature hepatocyte-like cells | SEES4 |

Human embryonic stem cell (hESC) lines SEES1, SEES4, and SEES5 were stably maintained in XF hESC culture medium containing 85% Knockout DMEM, 15% Knockout Serum Replacement XF CTS, 2 mM GlutaMAX-I, 0.1 mM NEAA, Pen-Strep, 50 µg/mL L-ascorbic acid 2-phosphate, 10 ng/mL heregulin-1β (recombinant human NRG-beta 1/HRG-beta 1 EGF domain), 200 ng/mL recombinant human IGF-1 (LONG R3-IGF-1; Sigma-Aldrich), and 20 ng/mL human bFGF (Akutsu et al., Regenerative Therapy; JCI insight). Undifferentiated hESCs were dissociated using dispase and plated on a dish coated with 0.1% human recombinant type I collagen peptide in 90 mm culture dishes. For hepatic differentiation, hESCs were cultured in XF hESC medium without growth factors (XF-KSR(-)) for 1 day and then in XF-KSR medium, which was replaced after 3 days with the XF hESC medium used as the differentiation medium. The differentiation medium was gently changed every 3–4 days until the indicated day.
